# BET inhibition sensitizes immunologically-cold Rb-deficient prostate cancer to immune checkpoint blockade

**DOI:** 10.1101/2022.03.24.485685

**Authors:** Brian Olson, Kiranj Chaudagar, Riyue Bao, Sweta Sharma Saha, Christina Hong, Srikrishnan Rameshbabu, Raymond Chen, Alison Thomas, Akash Patnaik

**Author notes:** **Financial Support:** This work was supported by the NCI Prostate SPORE Northwestern/University of Chicago (P50CA180995, A. Patnaik), Prostate Cancer Foundation Stephen A. Schwarzman PCF Young Investigator Award (B.M. Olson), Prostate Cancer Foundation Challenge Award (A. Patnaik and B.M. Olson), American Cancer Society Institutional Research Grant (Emory University, B.M. Olson), the Dunwoody Golf Club Prostate Cancer Research Award (a philanthropic award provided by the Winship Cancer Institute of Emory University, B.M. Olson), and Department of Defense Prostate Cancer Research Program Idea Development Award (W81XWH1910840, B.M. Olson). This research project was supported in part by the University of Chicago Center for Research Informatics Bioinformatics Core. This study was supported in part Institutes of Health under Award Number UL1TR002378. Research reported in this publication was also supported in part by the Pediatrics/Winship Flow Cytometry Core of Winship Cancer Institute of Emory University, Children’s Healthcare of Atlanta, the Emory Integrated Genomics Core (EIGC) shared resource of Winship Cancer Institute of Emory University, and NIH/NCI under award number P30CA138292. The content is solely the responsibility of the authors and does not necessarily represent the official views of the National Institutes of Health. To whom correspondence should be addressed: Brian Olson, Suite C3005, 1365 Clifton Road NE, Atlanta, GA 30322. To whom correspondence should be addressed: Akash Patnaik, KCBD 7152, 900 East 57th Street, Chicago, IL 60637. **Conflicts of Interest:** The authors declare no potential conflicts of interest.

## Abstract

**Purpose:** Non-T cell-inflamed immunologically “cold” tumor microenvironments (TME) are associated with poor responsiveness to immune checkpoint blockade (ICB), and can be sculpted by tumor cell genomics. Here we evaluated how Retinoblastoma (Rb) tumor suppressor loss of function (LOF), one of the most frequent alterations in human cancer and associated with lineage plasticity, poor prognosis and therapeutic outcomes, alters the TME, and whether therapeutic strategies targeting the molecular consequences of Rb loss enhance ICB efficacy.

**Experimental Design:** We performed bioinformatics analysis to elucidate the impact of endogenous Rb LOF on the immune TME in human primary and metastatic tumors. Next, we utilized isogenic murine models of Rb-deficient prostate cancer (PC) for *in vitro* and *in vivo* mechanistic studies to examine how Rb loss and bromodomain and extraterminal (BET) domain inhibition (BETi) reprograms the immune landscape, and evaluated *in vivo* therapeutic efficacy of BETi, singly and in combination with ICB and androgen deprivation therapy.

**Results:** Rb loss was enriched in non-T cell-inflamed tumors, and Rb-deficient murine tumors demonstrated decreased immune infiltration *in vivo*. The BETi JQ1 increased immune infiltration into the TME through enhanced tumor cell STING/NF-κB activation and type I interferon (IFN) signaling within tumor cells, resulting in differential macrophage and T cell-mediated tumor growth inhibition and sensitization of Rb-deficient PC to ICB.

**Conclusions:** BETi can reprogram the immunologically cold Rb-deficient TME via STING/NF-κB/IFN signaling to sensitize Rb-deficient PC to ICB. These data provide the mechanistic rationale to test combinations of BETi and ICB in clinical trials of Rb-deficient PC.

**STATEMENT OF TRANSLATIONAL RELEVANCE:** Rb LOF is one of the most common genomic alterations in human cancer, occurring in approximately 1/3 of advanced malignancies, Furthermore, loss of Rb correlates with enhanced aggressiveness and poor therapeutic outcomes. In this study, we demonstrate that loss of Rb is also associated with an immunosuppressive tumor microenvironment and lack of responsiveness to immune checkpoint blockade (ICB). As a strategy to overcome Rb LOF induced immunosuppression, we have demonstrated that BETi treatment drives STING/NF-κB signaling and type I interferon production within tumor cells, resulting in immune-mediated tumor control in Rb-deficient PC, which is accentuated by the combination with ICB and ADT. These findings provide a roadmap for maximizing the clinical translation of BET inhibitors into the clinic to treat aggressive-variant Rb-deficient PC.

## INTRODUCTION

Immune checkpoint blockade (ICB) has revolutionized cancer treatment, with the approval of several agents targeting CTLA4 and PD1/PD-L1 across dozens of malignancies. While approximately 20% of patients will generate robust, often durable, clinical responses, the majority of patients still do not benefit from ICB. Critically, the mechanisms to explain why this large subset of patients fail to respond are incompletely understood. In particular, metastatic, castrate-resistant prostate cancer (mCRPC) has shown limited responses to ICB, and the mechanisms for poor clinical activity in this disease remain inconclusive.

In a variety of malignancies, increased frequency of tumor-infiltrating lymphocytes (TILs) within the tumor microenvironment (TME) correlates with enhanced prognosis and responsiveness to immunotherapy (1). We have previously defined a ‘T-cell inflamed gene signature’ that includes markers of T cell infiltration, a select chemokine expression profile, and interferon (IFN) signaling (2), which predicts for enhanced responsiveness to ICB (3, 4). Given the association between this T-cell inflamed gene signature and responsiveness to ICB, the identification of mechanisms by which tumor cell intrinsic genomic alterations drive an inflamed or non-inflamed phenotype can identify potential predictive biomarkers and/or actionable targets for clinical response or resistance to immunotherapy.

One of the most common pathways altered in primary and metastatic cancers is silencing of the retinoblastoma (Rb) tumor suppressor signaling pathway (5). The canonical function of Rb is to regulate cell cycle progression in part through its interactions with transcription factor E2F, leading tumors to commonly disrupt Rb function to promote cancer development and progression (6). Rb is silenced (via loss of *Rb1* or its upstream regulator *Cdkn2a*) in nearly 1/3 of all metastatic patients, representing two of the four most common tumor suppressor alterations across metastatic disease (5). Rb loss-of-function (LOF) is also relevant in prostate cancer, being lost in a third of primary tumors (7), up to 75% of mCRPC patients (8), 90% neuroendocrine prostate cancer (9), and correlates with lineage plasticity (10) and poor clinical outcomes in metastatic disease (8). In addition to its canonical activity, the wide range of transcription factors that interact with Rb result in diverse cellular consequences within the TME following Rb LOF, including evasion of anti-tumor immunity (11). This can occur via decreased MHC II expression (12), increased PD-L1 expression (13), and modulation of a wide variety of genes associated with immune function (11,14,15). These data illustrate the broad tumor cell-intrinsic and -extrinsic effects of Rb loss within the TME, and suggests targeting the molecular consequences of Rb loss may reverse these immunosuppressive effects.

One of the critical factors in regulating the activity of proteins downstream of Rb are the bromodomain and extraterminal domain (BET) family of chromatin readers. BET family members recognize acetylated residues and serve as transcriptional scaffolds to help regulate gene transcription (16). This includes BRD4, which is central to mediating the activity of E2F following Rb loss (17). Given their critical role in regulating gene transcription, there has been considerable drug developmental efforts focused on using BET inhibitors (BETi) to target tumor cell growth, including in prostate cancer (18). Additionally, while BET inhibitors have not been specifically evaluated in Rb-deficient cancers, there is mechanistic overlap with the broader class of histone deacetylase inhibitors (HDACi) (19, 20), which have demonstrated some efficacy in Rb-LOF malignancies (21, 22), thus providing a rationale for a more targeted BETi-based approach that may exert similar effects compared with pan-HDACi. BET inhibitors have also been shown to enhance anti-tumor immune responses, modulating tumor cell chemokine and checkpoint ligand expression, inducing immunogenic cell death, and enhancing the infiltration and function of T cells into the TME (23–25). This includes prostate cancer, where BETi modulated gene expression and increased the immunogenicity of tumor cells (26). However, while BET inhibitors have shown promise in preclinical models when combined with immunotherapeutic strategies (24–27), the underlying molecular mechanism(s) of BETi-induced immune reprogramming in genomically-driven subsets of cancer remain to be elucidated. Furthermore, early-stage clinical trials of BET inhibitors have not demonstrated dramatic single-agent activity in most solid tumors, illustrating the need to identify predictive biomarkers of BET inhibitor efficacy, alone or in combination with ICB.

In this study, we tested the hypothesis that Rb LOF drives a non-T cell inflamed TME in cancer, which can be reprogrammed by BET inhibition to enhance immune-responsiveness in Rb-deficient PC. We demonstrate that Rb LOF is associated with a non-T cell inflamed microenvironment across numerous malignancies, including primary and metastatic PC. Using an isogenic murine model of Rb-deficient PC, we show that Rb loss decreases global immune infiltration into the TME, while enhancing STING expression within tumor cells. Treatment of Rb-deficient murine PC with BETi leads to DNA damage-induced activation of STING and NF-κB signaling within tumor cells, enhanced type I IFN production and increased immune infiltration into the TME. These immune-mediated effects led to differential tumor control in Rb-deficient versus Rb-proficient PC, which was dependent on STING/NF-κB signaling. Critically, the *in vivo* anti-cancer responses with BETi in Rb-deficient PC were enhanced with ICB and concurrent androgen deprivation therapy (ADT). These data provide a mechanistic rationale for the evaluation of BETi in combination with ICB in aggressive-variant immunotherapy-refractory Rb-deficient malignancies, including PC.

## MATERIALS AND METHODS

### Bioinformatic analysis of T cell-inflamed gene signature and genetic variations

RNAseq expression data, copy number and somatic mutations were obtained for primary cancer cohorts (from The Cancer Genome Atlas, n=6153 total patients analyzed, release date 2/4/2015) and the MET500 database (from the University of Michigan cohort, n=483 total patients, release date 8/17/2017) as published (2, 28). The RNAseq expression data contained upper quartile-normalized and log2-transformed RNAseq by expectation maximization (RSEM) values (excluding genes expressed in less than 80% of patients). Whole exome sequencing data was used to determine somatic mutation and copy number alterations for each subject. Unsupervised hierarchical clustering was performed using K-mean equal to 12 and Euclidean distance metrics. T cell-inflamed or non-inflamed clusters were selected using ConsensusClusterPlus v.1.16.0 based on previously-identified T cell-signature transcripts (*CD8A, CCL2, CCL3, CCL4, CXCL9, CXCL10, ICOS, GZMK, IRF1, HLA-DMA, HLA-DMB, HLA-DOA, HLA-DOB*) (2). Genes differentially expressed in T cell-inflamed and non-inflamed groups were detected using ANOVA and filtered using a false discovery rate (FDR) q value < 0.01 and a fold change > 2.0. Canonical pathways significantly enriched in *RB1* or *CDKN2A* loss were identified by Ingenuity Pathway Analysis (IPA – Ingenuity Systems, http://www.ingenuity.com) based on experimental evidence from the Ingenuity Knowledge database (release date 11/2015).

### TIMER bioinformatic analysis

Gene expression data (log_2_ RSEM values) from primary cancers, along with adjacent normal tissues in the TCGA database, were utilized for the estimation of difference in expression profile of RB1 across cancer types using the Tumor IMmune Estimation Resource (TIMER) web-server (https://cistrome.shinyapps.io/timer/). The analysis was performed across various tumor histologies (abbreviations as above). The gene expression of RB1 was further correlated with the estimated abundance of immune infiltrates using the “Gene” module to generate scatter plots and estimate the partial Spearman’s Correlation coefficient after correction for tumor purity. Somatic Copy Number Alteration (SCNA) status of RB1 was used to classify patients into either deep deletion or normal diploid genomic status, as defined by GISTIC2.0 (29), and the abundance of B cells, CD8+ and CD4+ T cells, macrophage, neutrophils, or dendritic cells was estimated in the different cancer types.

### Cell culture, generation of isogenic Rb-deficient cells, and drug treatments

Myc-CaP cells (generously provided by Dr. Leigh Ellis, Cedars-Sinai, Los Angeles, CA), as well as human PC3 and DU-145 cell lines (American Type Culture Collection, Manassas, VA, and authenticated by short tandem repeat profiling), were cultured in high-glucose Dulbecco’s Modified Eagles Medium supplemented with 10% FBS (Gemini Bioproducts, West Sacramento, CA), 2% L-glutamine, and 1% PenStrep (Thermo Fisher Scientific), hereafter referred to as DMEM/FCS. All cell lines were routinely tested to confirm lack of Mycoplasma infection (MycoAlert, Lonza, Basel, Switzerland). To generate isogenic Rb-deficient cell lines (Myc-CaP^ΔRb^), Myc-CaP cells were transfected with pCas9-Rb1-CRISPR plasmid (Sigma-Aldrich, St. Louis, MO) using Lipofectamine 2000 (Thermo Fisher Scientific, Waltham, MA). After 24 hours, cells were sorted based on GFP expression, and single GFP+ cells were sorted into 96-well plates using a FACSAria II (Becton Dickenson Biosciences, Franklin Lakes, NJ) cytometer and expanded. Expanded clones were tested for Rb expression via Western blotting (antibody clones listed in Table S1, Cell Signaling Technologies, Danvers, MA). For *in vitro* drug treatments, cells were treated with between 100pM-10μM JQ-1 (SelleckChem, Houston, TX) alone or in combination with 1μM H-151 (SelleckChem), 5μM BMS-345541 (SelleckChem), 1μM IKK-16 (SelleckChem), or the appropriate vehicle controls for the time indicated. As a positive control for activation of STING signaling and IFN-β expression, a one-hour treatment with the STING agonist 5,6-dimethylxanthenone-4-acetic acid (DMXAA, 50 μg/mL, Sigma-Aldrich) was used.

### Cell viability and apoptosis assays

Cells were treated with vehicle or JQ-1 for up to 72 hours and evaluated for cell growth via trypan blue staining (normalizing to vehicle control). For cell death assays, Myc-CaP^WT^ or Myc-CaP^ΔRb^ cells were treated with vehicle or increasing doses of JQ-1 for 72 hours, and analyzed for apoptosis/necrosis using the APC Annexin V Apoptosis Detection Kit with Propidium Iodide (BioLegend, San Diego, CA). Cells were analyzed using a BD Symphony Flow Cytometer (BD Biosciences), and the frequency of dead (Annexin V+/PI+) and dying (Annexin V+/PI-) cells were quantified. LD_50_ values were calculated using Quest Graph software (30).

### Western blotting

RIPA buffer supplemented with protease and phosphatase inhibitor cocktails (Roche, Basel, Switzerland) were used to prepare protein lysates from cell lines and tumor cell preparations. For Western blotting, the antibodies used are detailed in Supplementary Table 1. For semi-quantification of Western blot images captured and processed using Adobe Photoshop, ImageJ (National Institutes of Health) software was used to quantify densitometry for individual bands and normalized to respective control bands. To calculate IC_50_ and IC_90_ values based on HEXIM expression, densitometry results were analyzed using Quest Graph software (30).

### T cell migration assays

Migration of T cells was performed using inserts for 24-well plates with 5μm pores (Corning Life Sciences). Naïve T cells isolated from FVB splenocytes (EasySep, StemCell Technologies, Cambridge, MA) were placed into the top of a Boyden Chamber in DMEM/FCS. The bottom chamber contained either media alone or supernatants from Myc-CaP^WT^ or Myc-CaP^ΔRb^ cells treated with media alone or JQ-1 (alone or combined with inhibitors). The bottom chamber was also supplemented with either vehicle or CXCL10 (500 ng/mL, BioLegend) to promote T cell migration. Transwells were incubated for four hours, and cells that transmigrated into the lower chamber were collected and counted via trypan blue dye exclusion. Assays were conducted in triplicate and normalized to input.

### NanoString analysis

RNA was purified from Myc-CaP^WT^ or Myc-CaP^ΔRb^ cells treated with vehicle or 1μM JQ-1 for 24 hours using the RNEasy kit (Qiagen, Hilden, Germany). For NanoString analysis, purified RNA was used for hybridization using the NanoString mouse PanCancer Immune Profiling panel (NanoString Technologies, Seattle, WA). Data analysis was performed using nSolver 4.0, MSigDB, and R software.

### qRT-PCR

RNA was isolated from Myc-CaP^WT^ or Myc-CaP^ΔRb^ cells cultured in vehicle or JQ-1 (RNeasy, Qiagen). cDNA was synthesized using the iScript cDNA synthesis kit (BioRad, Hercules, CA) according to the manufacturer’s instructions. The SsoFast EvaGreen Supermix kit (BioRad) was used for qRT-PCR using recommended cycling conditions. Primers used are detailed in Supplementary Table 2.

### IFN-β ELISA

Cells were treated with vehicle or 1μM JQ-1 for 36 hours (alone or in combination with H-151 or BMS-345541), and supernatants were collected and evaluated for IFN-β secretion by ELISA (LEGEND MAX mIFN-β ELISA, Biolegend) according to the manufacturer’s instructions. For *ex vivo* ELISAs, tumors were homogenized, red blood cells were lysed using ACK buffer, and cells were plated out at 0.5x10^6^ cells/mL. After 24 hours, cells were treated with appropriate compounds, and incubated 36 hours prior to collection of supernatants and processed for IFNβ secretion by ELISA.

### Amnis ImageStream Analysis

Cells were seeded at density of 3x10^5^ cells per well in 6-well plates, and after 24 hours treated with vehicle, JQ-1 (1μM) or DMXAA (50μg/mL) for one hour. Following incubation, cells were collected, fixed and permeabilized with BD Pharmingen Transcription Factor Phospho (TFP) buffer set (BD Biosciences) and BD Phosflow Perm Buffer III (BD biosciences) per manufacturer protocol. Permeabilized cells were incubated with primary antibody cocktail targeting pIRF3 or pNFkB (clones listed in Supplementary Table 1). Cells were washed twice in TFP Perm/Wash buffer, and then incubated with fluorescently-labeled secondary antibodies for 1 hour. Lastly, nuclei were stained using DAPI (Invitrogen/Thermo Fisher Scientific, Waltham, MA) as per manufacturer protocol. Stained cells were imaged using Amnis ImageStream Mark II (Luminex, Austin, TX) system. The nuclear localization of pIRF3 and NFkB was quantified using IDEAS software 6.0 (Luminex).

### In vivo mouse studies

The Institute of Animal Care and Use Committees (IACUC) at Emory University and the University of Chicago approved all mouse procedures. Wild-type male FVB mice were obtained from Jackson Labs, and were challenged subcutaneously with 10^6^ Myc-CaP^WT^ or Myc-CaP^ΔRb^ cells and allowed to establish to a tumor volume of 50mm^3^. For treatment studies, animals were treated with the following compounds: JQ-1 (50mg/kg/d, reconstituted in DMSO and diluted 1/10 in 10% (w/v) hydroxypropyl-β-cyclodextrin), H-151 (750nmol/d, reconstituted in DMSO and diluted 1/10 in PBS/10% Tween-80, delivered intra-peritoneally (i.p.)), BMS-345541 (25 mg/kg/d, reconstituted in H_2_O, pH to 7.0 and delivered via oral gavage), degarelix (0.625 mg/kg, delivered i.p.), anti-CD4 antibody and anti-CD8 antibodies (BioXCell, Lebanon, NH – 200μg/treatment, diluted in PBS and delivered i.p.), or clodronate liposomes (Standard Macrophage Depletion kit, Encapsula Nanosciences – delivered i.p. at recommended dose of 300 μl of clodronate-liposomal emulsion containing 18.4 mM concentration of clodronate). Tumors were measured at least thrice weekly, and volume was calculated using the formula (π/6) x long diameter x (short diameter)^2^. Euthanasia was performed for mice bearing tumor ulceration or volumetric endpoint per IACUC-approved protocols.

### Multiparameter flow cytometric profiling of immune cells

Tumor or splenic tissue was processed into cell suspensions via gentle mechanical dissociation and filtered through a 70μm filter into single-cell suspensions. For evaluation of tumor-infiltrating immune cells, cells were treated with Ig block (TruStain FcX (anti-mouse CD16/32) antibody, BioLegend), and then stained with one of two flow panels (antibodies detailed in Supplementary Table 1): a lymphoid immune panel (Ghost Dye Red 780, human CD8, CD3, CD4, CD8, CD19, CD45, CXCR5, CD137, PD-1, Tim3, CXCR5, and NKG2D) or a myeloid immune panel (live/dead, human CD8, CD11b, CD11c, CD14, CD24, CD45, CD103, F4/80, Ly6C, Ly6G, I-A/I-E, PD-L1, and CD206). After surface staining, fixation (Cytofix, BD Biosciences), and permeabilization (PermWash, BD Biosciences), lymphocytes were intracellularly stained with Foxp3 and Ki67 (or IgG controls), and myeloid cells were intracellularly stained with Arg1 (or IgG control). Cells were analyzed on LSRII (BD Biosciences), BD Symphony (BD Biosciences) or Cytek Aurora (Cytek Biosciences, Bethesda, MD) flow cytometers. Cells were gated based on size/nucleation, singlets, live events, human CD8+ negative (dump channel), CD45+, and individual populations were identified as follows: CD4+ T cells (CD3+CD4+), CD8+ T cells (CD3+CD8+), NK-T cells (CD3+NKG2D+), NK cells (CD3-NKG2D+), GrMDSC (CD11b+MHCII-Ly6C^lo^Ly6G+), MoMDSC (CD11b+MHCII-Ly6G-Ly6C^hi^), Macrophage (CD11b+Ly6C+F4/80+), Monocyte (CD11b+Ly6C+F4/80-), Neutrophils (CD11b+Ly6C^lo^Ly6G+), DC1 cells (Ly6C-MHCII+CD24+CD11b+), DC2 cells (Ly6C-MHCII+CD24+CD103+), TAM1 cells (Ly6C-MHCII+F4/80+CD11b+CD11c-), TAM2 cells (Ly6C-MHCII+F4/80+CD11c+CD206+Arg1+), and Eosinophils (MHCII-CD11b+CD11c-SSC^hi^).

### Statistical Methods

Graph preparation and statistical analyses of *in vitro* and *in vivo* experiments was performed with the GraphPad Prism software (GraphPad Software Inc, San Diego, CA). Statistical significance for assays was assessed using a two-sided Fisher’s exact test, Welch’s corrected unpaired t-test, paired t-test, Wilcoxin signed-rank test, or non-linear regression analysis of tumor curve fits, as indicated within figure legends.

## RESULTS

### Immunologically cold metastatic and primary malignancies are enriched for loss of *Rb1*

As a first step towards understanding the mechanistic basis for intrinsic resistance of advanced malignancies to ICB, we analyzed approximately 500 metastatic samples from the MET500 database (5) for the expression of a T cell-inflamed gene signature. This analysis revealed that only approximately 1/3 of tumors are T cell-inflamed and the remainder display an intermediate phenotype or non-T cell-inflamed phenotype (Fig. 1A). Next, we analyzed the association between Rb LOF and the T cell inflamed versus non-inflamed status of metastatic samples. Interestingly, we observed that non-T cell inflamed metastatic tumors across all histologies were significantly enriched for either deletion or LOF mutations in *Rb1* (Fig. 1B).

**Figure 1.**
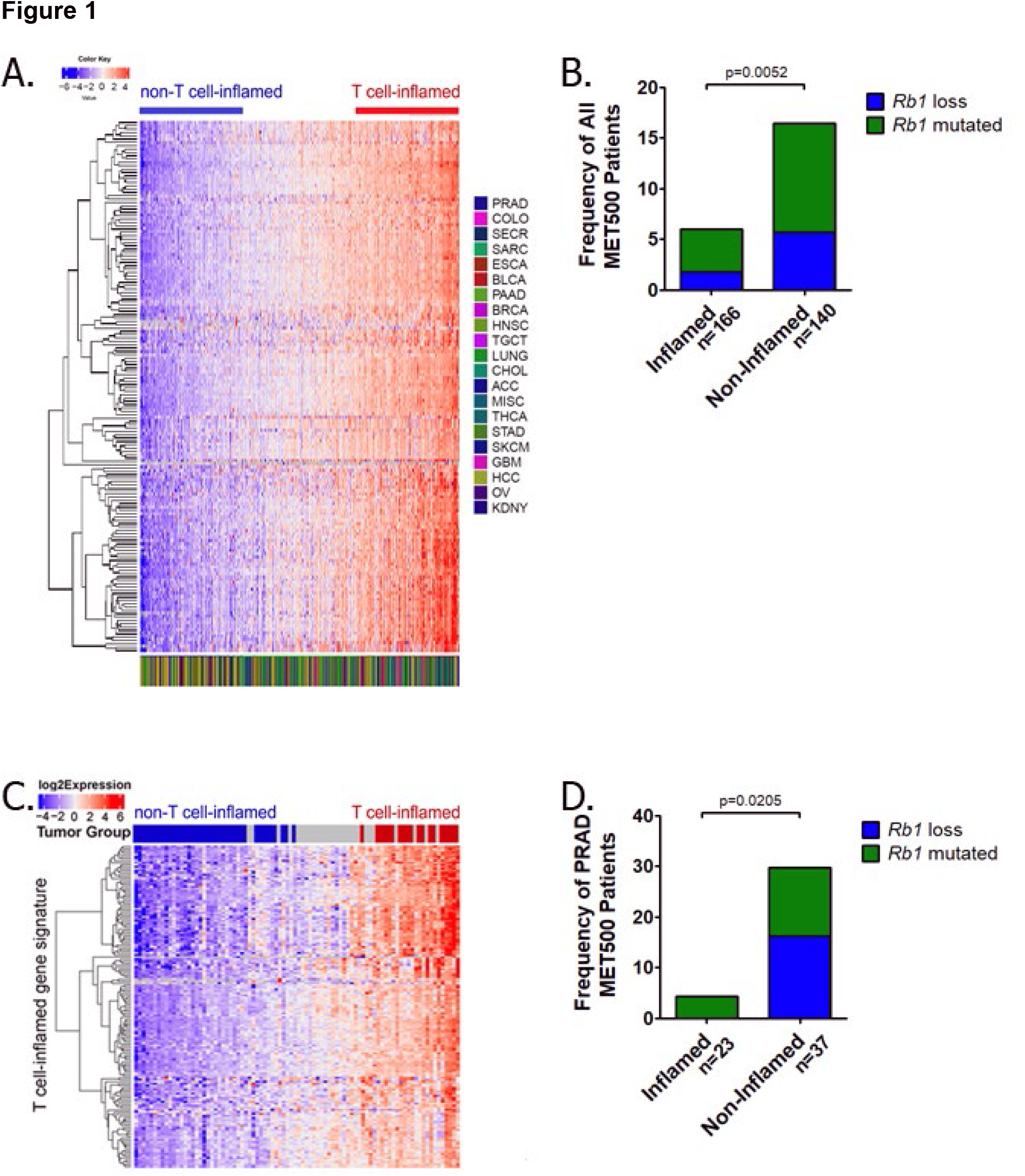
Loss of Rb is enriched in immunologically cold tumors. ***Panel A,*** Gene expression data from metastatic tumors contained in the TCGA MET500 database were analyzed for a validated expression panel of 13 genes that comprise a T cell-inflamed gene signature (2), along with other genes that cluster with this gene signature. Bar above heat map indicates tumors defined as T-cell inflamed (*red*) or non-T-cell inflamed (*blue*). The following abbreviations were used to indicate tumor types: ACC: Adrenocortical Carcinoma; BLCA: Bladder Urothelial Carcinoma; BRCA: Breast invasive carcinoma; CESC: Cervical squamous cell carcinoma; CHOL: Cholangiocarinoma; COAD: Colon Adenocarcinoma; DLBC: Lymphoid Neoplasm Diffuse Large B-cell Lymphoma; ESCA: Esophageal carcinoma; GBM: Glioblastoma multiforme; HNSC: Head and Neck squamous cell carcinoma; KICH: Kidney Chromophobe; KIRC: Kidney renal clear cell carcinoma; KIRP: Kidney renal papillary cell carcinoma; LAML: Acute Myeloid Leukemia; LGG: Brain Lower Grade Glioma; LIHC: Liver hepatocellular carcinoma; LUAD: Lung adenocarcinoma; LUSC: Lung squamous cell carcinoma; MESO: Mesothelioma; OV: Ovarian serous cystadenocarcinoma; PAAD: Pancreatic adenocarcinoma; PCPG: Pheochromocytoma and Paraganglioma; PRAD: Prostate adenocarcinoma; READ: Rectum adenocarcinoma; SARC: Sarcoma; SKCM: Skin Cutaneous Melanoma; STAD: Stomach adenocarcinoma; TGCT: Testicular Germ Cell Tumors; THCA: Thyroid carcinoma; THYM: Thymoma; UCEC: Uterine Corpus Endometrial Carcinoma; UCS: Uterine Carcinosarcoma; UVM: Uveal Melanoma. ***Panel B,*** metastatic tumors from the MET500 database were grouped into T cell-inflamed or non-T cell-inflamed tumors, and exomic and RNAseq data was used to quantify the frequency of patients with *Rb1* deletion or LOF mutations, as indicated in the panel. ***Panel C-D,*** Gene expression data from metastatic prostate tumors contained in the TCGA MET500 database (n=86 patients) were analyzed for a T cell-inflamed gene signature and other genes that cluster with this gene signature. The tumors were grouped into T cell-inflamed or non-T cell-inflamed tumors (C) and used to quantify the frequency of patients with *Rb1* deletion or LOF mutations (D).

Next we focused on mCRPC, which has demonstrated a general lack of responsiveness to ICB and in which Rb is lost in up to 75% of patients (8, 31). We observed a predominance of immunologically non-T cell inflamed tumors within mCRPC patients (Fig. 1C), which were enriched for *Rb1* LOF (Fig. 1D). Similar observations were found when analyzing several primary malignancies within the TCGA database, including primary PC, in which loss or LOF mutation of *Rb1* or its upstream regulator *Cdkn2a* was enriched in non-T cell inflamed primary tumors (Supplementary Fig. 1).

To further evaluate the effects of Rb expression on the tumor immune microenvironment, we utilized the TIMER predictive algorithm and observed decreased *Rb1* RNA expression in PC tissue compared to normal tissue (Fig. 2A). Furthermore, when the amplitude of *Rb1* expression was quantified as a function of CD8 infiltration, a positive correlation was identified between *Rb1* expression and CD8 infiltration (Fig. 2B). When the same data was analyzed by stratifying patients based on genomic deletion of *Rb1*, the results demonstrated that patients with biallelic deletion of *Rb1* had decreased infiltration of multiple immune populations, including CD4+ and CD8+ T cells as well as neutrophils (Fig. 2C). Decreased immune infiltration in patients with biallelic deletion of *Rb1* was similarly observed in 7/11 tumor histologies in which loss of *Rb1* and/or *Cdkn2a* was enriched in immunologically-cold non-T cell-inflamed tumors (Supplementary Fig. 2). Taken together, these data indicate that loss of Rb results in immunosuppressive non-T cell inflamed phenotypes across multiple cancer histologies.

**Figure 2.**
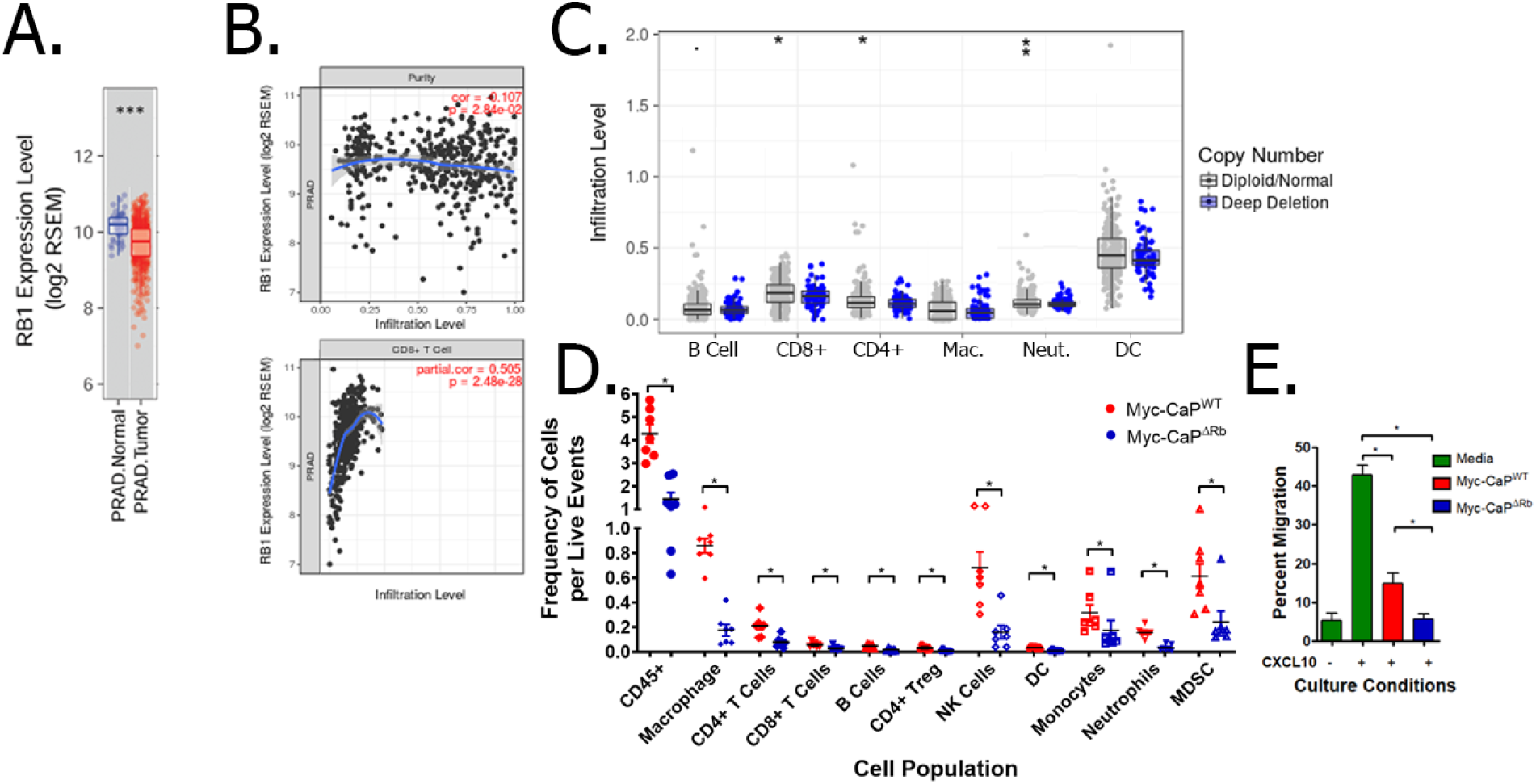
Rb-deficient prostate cancer has decreased immune infiltration in the tumor microenvironment. ***Panel A,*** Gene expression data from primary prostate cancer along with the adjacent normal tissues in the TCGA database. *** indicates p<0.001 by Wilcoxin signed-rank test. ***Panel B,*** Gene expression level of *Rb1* was compared with estimated abundance of tumor cells (to measure tumor purity; *top panel*) and CD8 T cells (*bottom panel*) using TIMER. ***Panel C,*** Somatic Copy Number Alteration (SCNA) data from the primary cancers in the TCGA database was utilized for the estimation of copy number status of *Rb1* in prostate tumors (either wild-type or deep deletion), and were analyzed for estimated B cell, CD4+ and CD8+ T cell, macrophage (Mac.), neutrophil (Neut.), and dendritic cell (DC) frequencies using TIMER. *: p < 0.05; **: p-value <0.01; ***: p<0.001, all by Wilcoxin signed-rank test. ***Panel D,*** Wild-type male FVB mice were challenged with Myc-CaP^WT^ or Myc-CaP^ΔRb^ tumor cells. Once tumors reached 200 mm^3^, tumors were collected and evaluated for frequency of various tumor-infiltrating immune populations within wild-type (*red*) or Rb-deficient (*blue*) tumors. The data is normalized to the number of live events. * indicates p<0.05 by Student’s t-test. ***Panel E,*** supernatants from MycCaP^WT^ or Myc-CaP^ΔRb^ tumor cells were evaluated for their ability to suppress CXCL10-induced T-cell migration across a transwell. * indicates p<0.05 by Student’s t-test.

### Targeted deletion of *Rb1* results in decreased immune infiltration into the prostate tumor microenvironment *in vivo* and an immunosuppressive phenotype in PC cells *in vitro*

As a first step towards developing a murine model of Rb-deficient PC for experimental therapeutics, we performed CRISPR-mediated deletion of *Rb1* in Myc-CaP cells (derived from a transgenic model of probasin-driven c-myc-overexpression in mouse prostate (32)). Interestingly, we observed no significant difference in *in vitro* and *in vivo* kinetics as a function of Rb loss (Supplementary Fig. 3). However, when FVB/NJ mice bearing syngeneic wild-type (Myc-CaP^WT^) or Rb-deficient (Myc-CaP^ΔRb^) tumors were evaluated for immune infiltration into the TME *in vivo*, Rb-deficient tumors were found to have decreased infiltration of several immune populations, including a 3-fold decrease in overall CD45+ cells, and a 2.6- and 2.1-fold decrease in CD4+ and CD8+ T cells, respectively (Fig. 2D). Additionally, supernatants from Myc-CaP^ΔRb^ cells were able to suppress T cell migration across a transwell to a greater extent than supernatants from Myc-CaP^WT^ cells (Fig. 2E). To elucidate the mechanistic basis for the observed immunosuppressive phenotype in Rb-deficient cancers, we performed differential transcriptomic analysis of Myc-CaP^WT^ and Myc-CaP^ΔRb^ cells *in vitro* (Supplementary Fig. 4). Interestingly, loss of *RB1* was found to result in alterations in gene expression that can promote an immunosuppressive tumor microenvironment. This included an increase in at least five genes associated with immunosuppression (including *Ido1)* and a decrease in 20 genes associated with immune stimulation (such as *Tnfsf10*, which is regulated by transcription factors controlled by Rb (11)). However, amongst the upregulated immunostimulatory genes, we found that loss of Rb was also associated with increased expression of *Tmem173* (a.k.a. stimulator of interferon genes, STING) at both the RNA (Supplementary Fig. 4A-B) and protein level (Supplementary Fig. 4C). Collectively, these *in silico*, *in vitro,* and *in vivo* data demonstrate that Rb LOF drives an immunosuppressive tumor microenvironment, but a paradoxically enhanced expression of STING within tumor cells.

### BET inhibition drives increased type I interferon expression in Rb-deficient tumor cells via enhanced DNA damage-induced non-canonical STING/NF-κB signaling

Given that Rb loss is associated with an immunosuppressive TME, identifying means of reversing the broad molecular consequences of Rb loss would have the potential of overcoming this immunosuppression, thereby enhancing efficacy of immunotherapy. Previous studies have identified a role for BET family proteins in mediating the activity of transcription factors that are regulated by Rb (17). Based on our findings that Rb LOF increases STING expression and prior work demonstrating that BET inhibitors can induce DNA damage (33), we hypothesized that BETi selectively increases type I interferon production in Myc-CAP^ΔRb^ cells via STING pathway activation. We first performed *in vitro* dose-pharmacodynamic evaluation of the pan-BET inhibitor JQ-1 (34), in Myc-CAP^WT^ and Myc-CAP^ΔRb^ cells, as assessed by increase in HEXIM-1 expression (Supplementary Figure 5). The observed IC_90_ concentrations of JQ-1 were found to be approximately 10-fold below the LD_50_ concentrations needed to directly mediate cell death (Supplementary Figure 6), permitting the evaluation of tumor cell-extrinsic mechanisms of action of BET inhibition in Rb-deficient disease. Treatment with sublethal IC_90_ concentrations of JQ-1 selectively increased IFN-β RNA and protein expression in Myc-CAP^ΔRb^ cells, relative to wild-type cells (Figure 3A). To demonstrate the mechanistic basis for this observation, we analyzed DNA damage-induced STING signaling in Myc-CAP^WT^ and Myc-CAP^ΔRb^ cells following JQ1 treatment. We observed that JQ-1 increased γ-H2AX expression, a marker of DNA double-strand breaks (DSB), in both wild-type and Rb-deficient tumor cells (Fig. 3B). Furthermore, while STING expression was increased at baseline in Myc-CaP^ΔRb^ tumor cells, JQ-1 did not lead to increased phosphorylation of TBK1 or IRF3, the canonical signaling pathway associated with increased type I interferon expression. Interestingly, we observed a time-dependent increase in phosphorylated NF-κB specifically in Myc-CaP^ΔRb^ cells (Fig. 3B), a non-canonical STING-induced signaling pathway that can also drive increased type I interferon expression and immune infiltration. To validate the role of NF-κB versus IRF3 signaling, imaging flow cytometry demonstrated increased nuclear translocation of phosphorylated NF-κB (not phosphorylated IRF3) following JQ-1 stimulation selectively in Myc-CaP^ΔRb^ cells (Fig. 3C-D and supplementary Fig. 7). Critically, concomitant inhibition of STING with H-151 (STING inhibitor) or NF-κB with BMS-345541 or IKK-16 (I kappa B kinase inhibitors that block NF-kappa B-dependent transcription) abrogated JQ-1-induced IFN-β expression by Myc-CaP^ΔRb^ cells *in vitro* (Fig. 3E) and homogenized Myc-CaP^ΔRb^ tumor-derived single cell suspensions *ex vivo* (Fig. 3F). Additionally, treatment with BET inhibition restored T cell migration towards Myc-CaP^ΔRb^ cells in a STING- and NF-κB-dependent fashion (Fig. 3G). These data demonstrate that BET inhibitors selectively activate the STING/non-canonical NF-κB pathway to induce type I IFN signaling in Rb-deficient prostate cancer, relative to its Rb-proficient counterpart.

**Figure 3.**
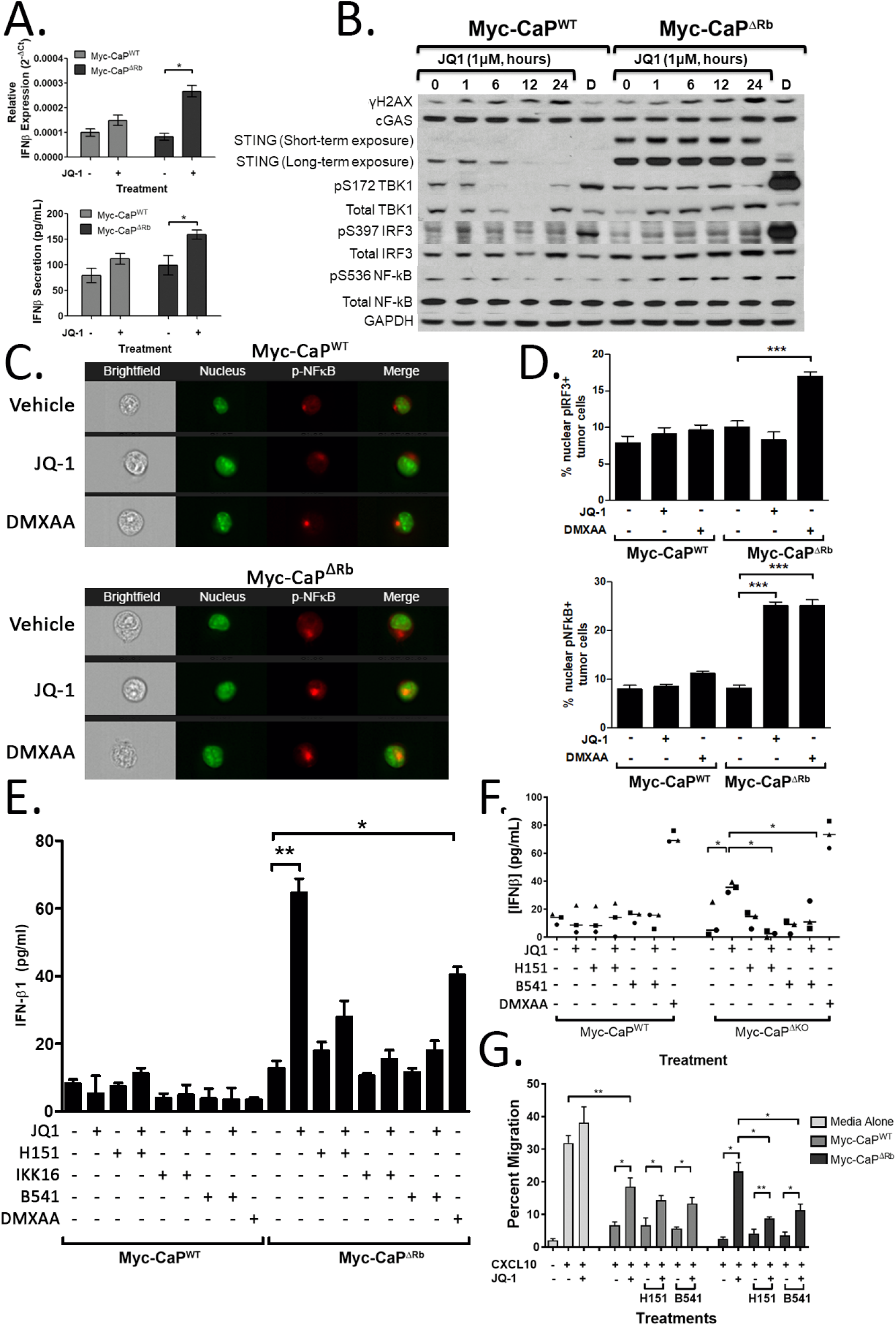
BETi induces DNA damage and type I IFN expression through NFκB signaling. ***Panel A***, Expression of IFNβ was measured at the RNA level by qRT-PCR (*top panel*) or at the protein level by ELISA (*bottom panel*) using RNA or supernatants from Myc-CaP^WT^ and Myc-CaP^ΔRb^ cells treated with vehicle or 1μM JQ-1. ***Panel B,*** Myc-CaP^WT^ or Myc-CaP^ΔRb^ cells were treated with vehicle or 1μM JQ-1 for 1-24 hours (or a DMXAA positive control ‘D’, 50μg/mL for one hour) and evaluated for γH2AX, c-GAS, STING, phosphorylated and total IRF3, phosphorylated and total TBK1, phosphorylated and total NF-κB, and GAPDH expression by Western blot. Low and high exposure for STING expression is provided. ***Panel C,*** Myc-CaP^WT^ *(top panels)* or Myc-CaP^ΔRb^ (*bottom panels)* cells were treated with 1μM JQ-1 (or vehicle or DMXAA controls) for one hour, and nuclear expression of pNF-κB was measured via ImageStream analysis. Panels demonstrate brightfield images, nuclear staining (green), p-NF-κB (red), or merged images. ***Panel D***, ImageStream data was quantified to identify the frequency of nuclear translocation of p-IRF3 (*top panels*) or p-NF-κB (*bottom panels*) in vehicle, JQ-1, or DMXAA-treated Myc-CaP^WT^ or Myc-CaP^ΔRb^ cells. ***Panel E***, Myc-CaP^WT^ or Myc-CaP^ΔRb^ cells were treated with 1μM JQ-1 (or vehicle or DMXAA controls) alone or in the presence of the STING inhibitor H-151 or the NF-κB signaling inhibitors IKK-16 or BMS-345541 (*B541*). Supernatants were collected and measured for IFN-β secretion via ELISA. ***Panel F***, untreated tumors from animals bearing Myc-CaP^WT^ or Myc-CaP^ΔRb^ tumors were processed into a single cell suspension, and then treated with 1μM JQ-1 or controls alone or in combination with H-151 or BMS-345541 (or vehicle or DMXAA controls) for 36 hours. Supernatants were collected and evaluated for IFN-β secretion by ELISA. Each dot represents the average of three experimental replicates from tumor cells isolated from one tumor-bearing mouse, with a total of three wild-type or Rb-deficient tumor-bearing mice used for biological replicates. * indicates p<0.05 by paired student’s t-test. ***Panel G***, supernatants from Myc-CaP^WT^ or Myc-CaP^ΔRb^ cell lines treated with vehicle or 1μM JQ-1 alone or in combination with vehicle, H-151, or BMS-345541 for 24 hours were evaluated for their ability to suppress CXCL10-induced T cell migration across a transwell *in vitro*. * indicates p<0.05 by student’s t-test.

### BET inhibition elicits STING/NF-κB-dependent and T cell/macrophage-dependent anti-cancer immune responses, resulting in selective tumor growth delay in mice bearing Rb-deficient tumors *in vivo*

Based on the observation that BET inhibition promotes an inflammatory immune phenotype via selective DNA damage-induced STING signaling and type I interferon expression within Rb-deficient cancer cells *in vitro*, we next tested the possibility that *in vivo* treatment with JQ1 reverses the immunosuppressive TME in the syngeneic Rb-deficient murine PC model *in vivo*. Treatment with JQ-1 for seven days led to an increase in DNA DSB in mice bearing both Myc-CAP and Myc-CAP^ΔRb^ tumors (Fig. 4A), but a significant increase in IFN-β expression within the TME only in the Myc-CAP^ΔRb^ tumors (Fig. 4B). Furthermore, treatment with BET inhibition significantly increased infiltration of both effector and myeloid immune populations, particularly in Myc-CaP^ΔRb^ tumors. This included an approximately 4-fold and 3-fold increase in CD4+ and CD8+ T cells, respectively (Fig. 4C). In contrast, BETi treatment increased TAM2 immunosuppressive macrophages in wild-type tumors, not observed in Myc-CAP^ΔRb^ tumors (Fig. 4C). Additionally, concurrent treatment of Myc-CaP^ΔRb^ tumor-bearing animals with JQ-1 and H-151 or BMS-345541 resulted in an approximately 3.5-fold and 6-fold decrease in T cell infiltration, relative to control mice (Fig. 4D). These *in vivo* data illustrate that non-canonical STING/NF-κB signaling plays a central role in mediating BETi-induced immune infiltration into the immunologically-cold Rb-deficient prostate tumor microenvironment.

**Figure 4.**
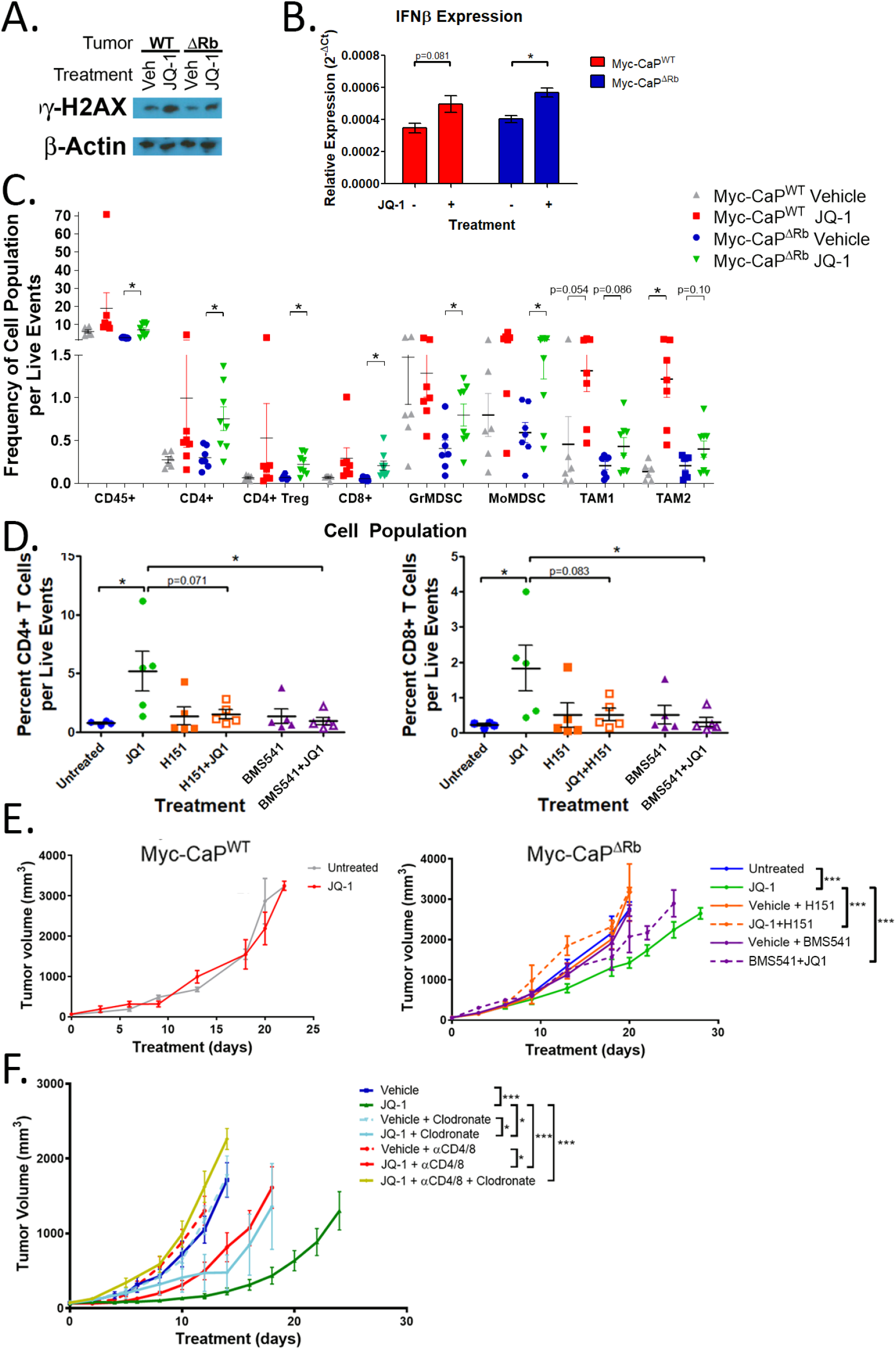
BET inhibition increases immune infiltration and delays tumor growth in immunosuppressive Rb-deficient tumors in a STING/NFκB- and T cell/macrophage-dependent fashion. Wild-type FVB mice were challenged with Myc-CaP^WT^ or Myc-CaP^ΔRb^ cells, and tumors were allowed to establish to 50mm^3^, and then treated for seven days with vehicle or JQ-1 (50 mg/kg/d). ***Panel A***, protein lysates from harvested tumors were evaluated for expression of γH2AX or β-actin by Western blot. ***Panel B***, RNA isolated from tumors were evaluated for expression of IFN-β by qRT-PCR. * indicates p<0.05 by student’s t-test. ***Panel C,*** Tumors were harvested and analyzed for immune infiltration by flow cytometry. * indicates p<0.05 by student’s t-test. ***Panel D***, Myc-CaP^ΔRb^ tumor-bearing animals were treated for seven days with vehicle, JQ-1, H-151, or BMS-345541, or JQ1 combined with either H-151 or BMS-345541. Following seven days of treatment, tumors were collected and evaluated for CD4+ (*left panel*) or CD8+ (*right panel*) infiltration via flow cytometry. ***Panel E***, Wild-type FVB mice from both Myc-CaP^WT^ (*left panel*) or Myc-CaP^ΔRb^ (*right panel*) groups were treated with vehicle or JQ-1, and animals bearing Myc-CaP^ΔRb^ tumors were also treated with either H-151, BMS-345541, or JQ1 combined with H-151 or BMS-345541. Animals were treated daily and followed for tumor growth until tumors reached 3000 mm^3^. * indicates p<0.0001 by non-linear regression analysis of tumor curve fits. ***Panel F***, Mice bearing ∼50mm^3^ Myc-CaP^ΔRb^ tumors received either CD4/CD8 depletion antibodies (on days -3, -1, 3, 7 and then weekly), macrophage-depleting clodronate liposomes (day -1 and then weekly), or control until the completion of the study. The experimental groups included mice receiving vehicle, JQ-1 or immune depleting treatments alone, singly and in combination as indicated. * indicates p<0.01 and *** indicates p<0.0001 by non-linear regression analysis of tumor curve fits.

To determine whether JQ-1-driven anti-cancer immune responses control long-term tumor growth, animals bearing wild-type or Rb-deficient tumors were treated daily with vehicle or JQ-1, alone or in combination with STING or NF-κB inhibition. Treatment with JQ-1 was found to delay tumor growth in animals bearing Myc-CaP^ΔRb^ tumors, but not in wild-type tumor-bearing animals (Fig. 4E). However, concomitant treatment of JQ-1 with H-151 or BMS-345541 resulted in abrogation of the anti-cancer activity observed with JQ-1 alone (Fig. 4E).

To evaluate the relative contribution of tumor cell intrinsic versus tumor cell extrinsic mechanisms on JQ-1-mediated tumor control, depletion of specific immune cell subsets were conducted. Mice bearing Rb-deficient tumors received JQ-1 alone or in combination with macrophage or CD4/CD8 T-cell depletion (Supplementary Fig. 8). While JQ-1 was found to delay tumor growth in Myc-CaP^ΔRb^ knockout tumor bearing mice, this anti-tumor benefit was partially abrogated with macrophage (p<0.01) or T cell (p<0.0001) depletion, and completely abolished with dual T cell/macrophage depletion (Figure 4F). Collectively, these data demonstrate that JQ-1 induces macrophage and T cell-mediated tumor control via non-canonical STING/NF-κB signaling in Rb-deficient prostate cancer.

### PD-L1 blockade and ADT enhance the anti-tumor efficacy of BET inhibition

While BET inhibition increased infiltration of effector immune populations into the TME, we also observed an increase in suppressive immune populations such as Tregs and myeloid-derived suppressor cells (Fig. 4C). Furthermore, tumor-infiltrating T cells showed a trend towards increased PD-1 expression relative to vehicle controls. Importantly, TAM2 and Gr-MDSC myeloid populations showed a statistically significant increase in PD-L1 expression in JQ-1-treated tumor-bearing mice, relative to vehicle controls (Supplementary Fig. 9). These data demonstrate the relevance of the PD-1/PD-L1 signaling axis in Rb-deficient tumors treated with JQ-1, and suggested combining BETi with blockade of this axis may enhance anti-tumor responses. To this end, mice bearing Rb-deficient tumors were treated with either vehicle or JQ-1 alone or combined with anti-PD-L1 and followed for tumor growth. As demonstrated previously, treatment with JQ-1 alone led to a significant delay in tumor progression in Rb-deficient tumor bearing animals compared to treatment with JQ-1 alone (Fig. 5A). However, this was significantly enhanced when combined with anti-PD-L1 blockade, and not observed with anti-PD-L1 blockade alone (Fig. 5A).

**Figure 5.**
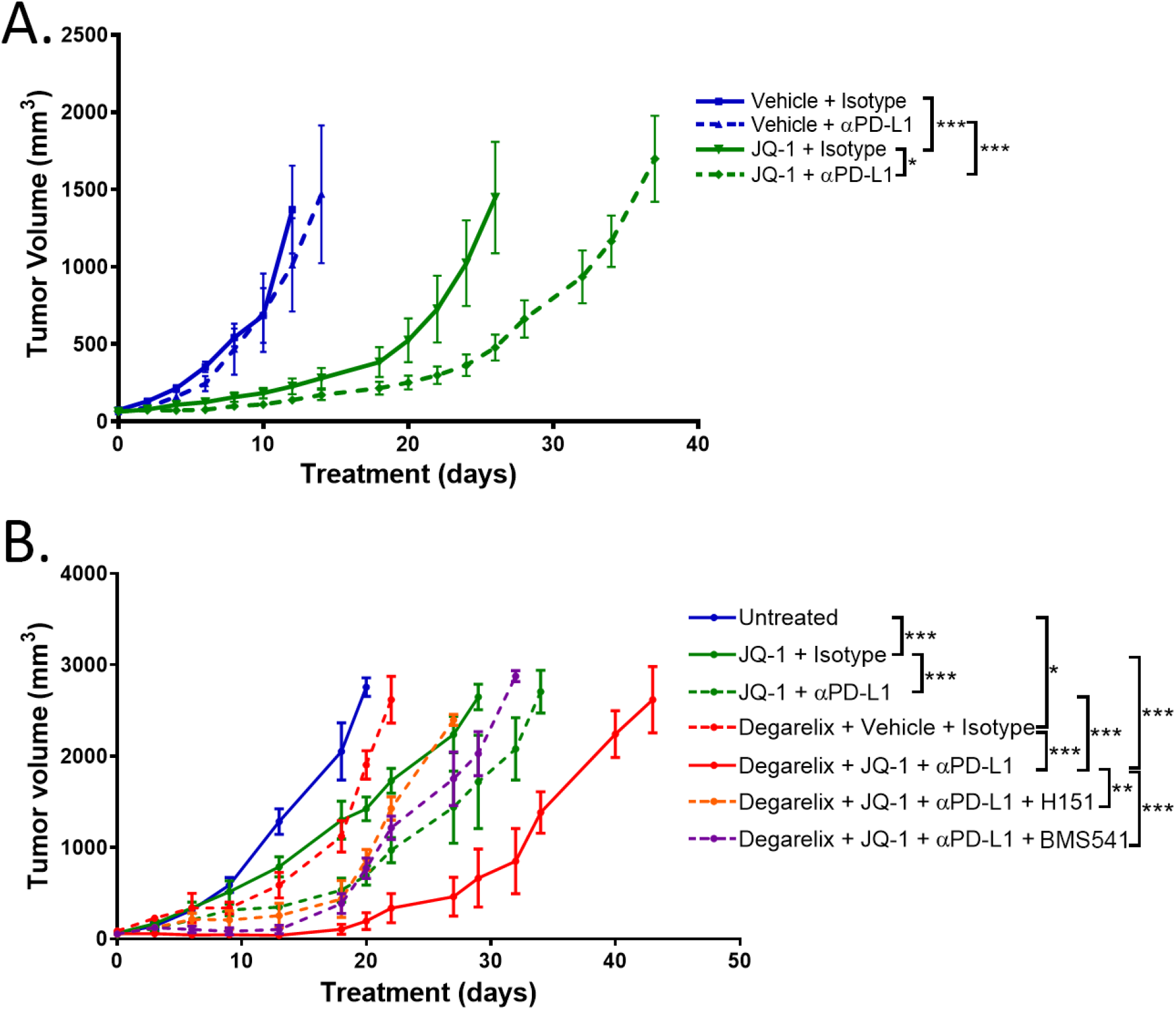
PD-L1 blockade enhances the anti-tumor activity of BET inhibition alone or in combination with ADT in Rb-deficient prostate cancer. ***Panel A***, Myc-CaP^ΔRb^ tumor-bearing animals were treated daily with vehicle or JQ-1 and either anti-PD-L1 or IgG control on days 7, 10, 14, 21, and 28 following the initiation of vehicle or JQ-1 treatment and followed for tumor growth. *: p<0.01; *** : p<0.0001 via nonlinear regression analysis of tumor growth curves. ***Panel B,*** Myc-CaP^ΔRb^ tumor-bearing mice were divided into groups receiving degarelix or vehicle. Vehicle-treated animals received JQ-1 alone (*solid green*) or JQ-1 with PD-L1 (*dashed green*), as in panel A. Degarelix-treated animals received vehicle or isotype controls (*dashed red*), JQ-1 and PD-L1 (*solid red*), or JQ-1 and PD-L1 combined with daily H-151 (*dashed orange*) or BMS-345451 (*dashed purple*). *: p<0.05; **: p<0.001; *** : p<0.0001 via nonlinear regression analysis of tumor growth curves.

**Figure 6.**
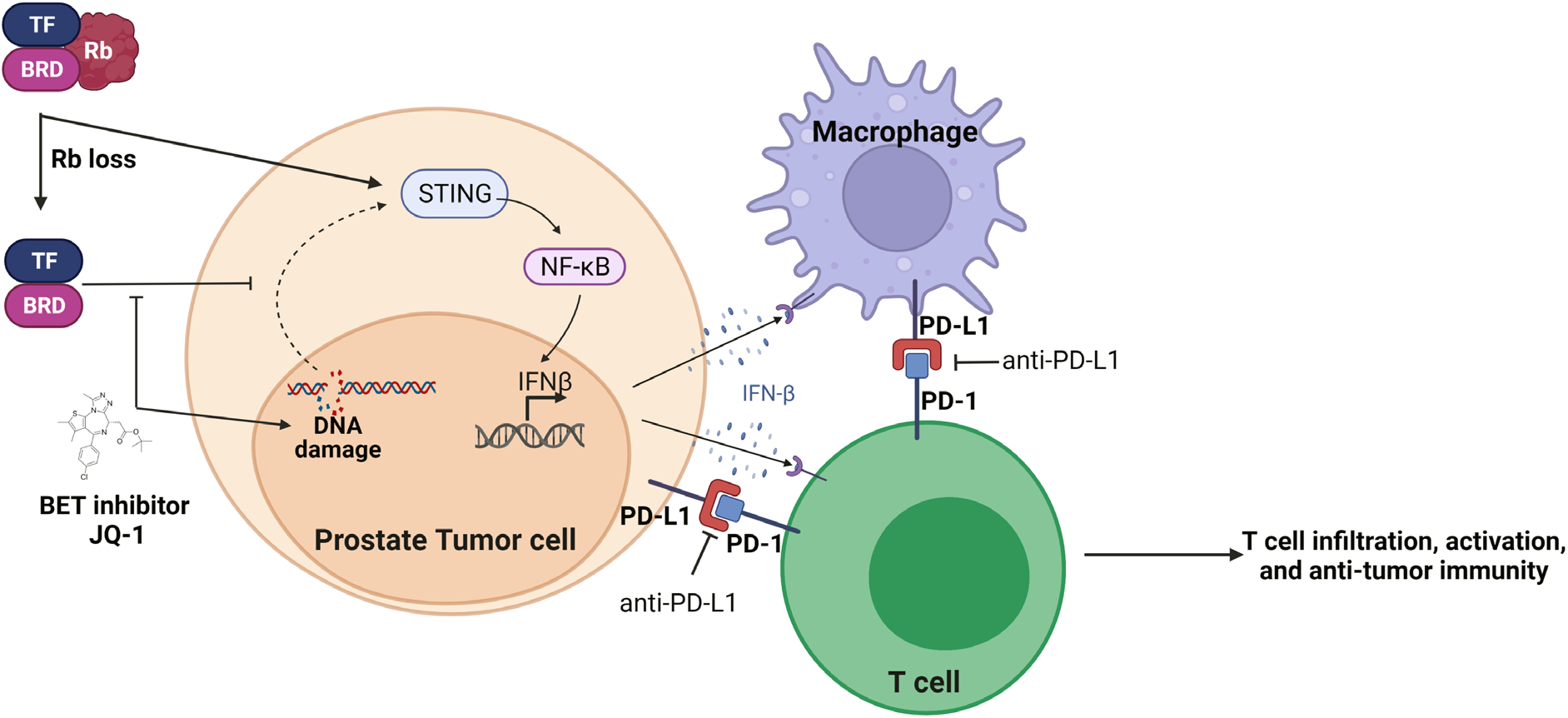
Mechanistic model for BET inhibitor-mediated sensitization to ICB in Rb-deficient PC. When Rb is lost in prostate tumor cells, Rb-regulated transcription factors (such as E2F, GATA, and others) are released and allowed to mediate gene transcription that is dependent on BET family of proteins. This altered transcriptional program promotes an immunosuppressive phenotype in PC cells that can suppress T cell migration *in vitro* and *in vivo*. However, Rb loss also results in upregulation of STING, which sensitizes these tumor cells to DNA damage signaling induced by BET inhibition. BETi activates STING and non-canonical NF-κB signaling, type I interferon secretion, and a macrophage and T cell dependent anti-cancer immune response, which is enhanced by ICB and/or ADT.

To further potentiate the anti-tumor immune responses mediated by JQ-1, Rb-deficient tumor bearing mice were treated also with androgen deprivation therapy (ADT) using degarelix (chemical castration), a standard-of-care treatment for advanced prostate cancer that we and others have demonstrated elicits increased immune infiltration in wild-type Myc-CaP tumor-bearing mice (35). Consistent with prior clinical data (36), while degarelix alone did not induce signficant anti-tumor responses in Rb-deficient tumor-bearing animals, the addition of degarelix to JQ-1 and anti-PD-L1 significantly enhanced the anti-tumor response elicited by corresponding singlet and doublet therapies (Fig. 5B). This enhanced responsiveness was abolished with concurrent H-151 or BMS-345541, further demonstrating the critical role of STING and NF-κB signaling in mediating the anti-tumor efficacy of ADT/BETi/aPD-L1 triple combination therapy in Rb-deficient PC.

## DISCUSSION

The prevalence and functional consequences of Rb loss across tumors of all histologies represents a significant clinical challenge, and the development of personalized therapeutic approaches targeting this genomic aberration has proved elusive. Silencing of the Rb pathway occurs in at least 1/3 of patients with metastatic disease, either through mutually exclusive LOF mutations of *Rb1* or its upstream regulator *Cdkn2a* (5). Furthermore, Rb loss often occurs in advanced disease, particularly associated with lineage plasticity (10) and neuroendocrine differentiation (9), and predicts for poor clinical outcomes (8). However, targeting the consequences of Rb loss have proved challenging; while targeting the activity of specific pathways deregulated by Rb LOF has demonstrated a direct cytotoxic effect against Rb-deficient tumor cells in preclinical models, including checkpoint kinase (CHK) inhibitors in a preclinical model of triple-negative breast cancer (37), the pleiotropic effects of Rb loss ultimately render Rb-deficient cancers resistant to these approaches. Conversely, the utilization of therapeutic strategies aimed at more broadly targeting the molecular consequences of Rb LOF (such as histone deacetylase or DNA methyltransferase inhibitors), face the challenge of significant toxicities when translated into the clinic (38).

The discovery and therapeutic targeting of cross-talk mechanisms between tumor cell-intrinsic genomic alterations and the immune microenvironment will enable identification of rational combinatorial strategies to enhance ICB-responsiveness in immune-refractory cancers. Emerging evidence across several diseases has demonstrated that genomic alterations in tumor cells facilitate the evasion of anti-tumor immune responses. This includes activation of the WNT/β-catenin and PTEN pathways in metastatic melanoma and several other cancers (2,28,39). Here we show that loss of Rb across multiple primary and metastatic histologies is associated with a non-T cell inflamed gene signature. Utilizing a reverse-translation approach in a syngeneic model of Rb-deficient PC, we show loss of Rb results in decreased immune infiltration into the TME. Furthermore, treatment of Rb-deficient PC with BETi induces DNA DSB sensing STING/NF-κB activation and downstream type I IFN expression, resulting in enhanced single agent efficacy of BETi *in vivo*, relative to isogenic Rb-proficient PC, and sensitization to PDL1 blockade. Importantly, we demonstrate that the non-tumor cell autonomous effects of BETi are the predominant driver of anti-tumor responsiveness in Rb-deficient PC, as the anti-tumor benefit following BET inhibition was dependent on T cell and macrophage activity. Collectively, these data provide a scientific rationale for evaluation of single-agent BETi in Rb-deficient disease, as well as in combination clinical trials of BETi with ICB in Rb-deficient patients, who may otherwise fail to respond to immunotherapy.

Loss of *Rb1* acts as a ‘first-hit’ for BETi-induced DNA damage, as Rb-deficient cancer cells have diminished DNA repair activity due in part to slowing of the replication fork progression during S-phase, thereby enhancing susceptibility to DNA damage (40). This slowing of the replication fork is also linked to the activity of BET inhibitors in promoting DNA DSB and NF-κB signaling (33, 41). However, this activity of BETi would be predicted to be maximized when cells have already received a first hit, as in BRCA-deficient breast cancer (42, 43). This provides a rationale for testing these agents in Rb-deficient disease, in which Rb loss increases tumor cell susceptibility to DNA damage, and lays the groundwork for future studies aimed at combining BETi (or alternative means of inducing DNA damage and STING signaling (37)) with immunotherapy in Rb-deficient cancers.

While cGAS/STING pathway activation is mediated through canonical TBK1/IRF3 signaling, non-canonical NF-κB signaling is also a well-established means of inducing type I IFN. The presence of cytosolic DNA DSB allosterically activate cGAS, resulting in the generation of cyclic dinucleotide cGAMP, STING activation and subsequent TBK1 and IRF3 phosphorylation and type I IFN production. However, in the context of nuclear DNA damage, STING is activated and leads predominantly to NF-κB activation and downstream type I interferon expression (44). Since the use of BET inhibitors in Rb-deficient disease has the potential of inducing nuclear DNA DSB, we hypothesized that BETi-induced DNA damage activates NF-κB, as opposed to TBK1/IRF3 signaling. While the anti-tumor activity of BETi in a murine melanoma model was previously linked to the activity of NF-κB, this was in the context of tumor cell-intrinsic suppression of the oncogenic driver Spp1 (45). Alternatively, in a murine model of triple-negative breast cancer, NF-κB signaling in tumor-associated macrophages (TAM) promoted resistance to BETi (46). These reports complement the novelty of our findings, where BETi activates NF-κB signaling in Rb-deficient tumor cells harbouring intrinsic susceptibility to DNA damage, resulting in a macrophage and T cell-dependent tumor cell-extrinsic anti-tumor immune response.

As a consequence of Rb loss, BET family members are known to regulate the activity of several transcription factors particularly relevant in prostate cancer. For example, increased E2F transcriptional regulation is mediated through BRD4, which can play a role in regulating the transcriptional activity of the androgen receptor (AR) (47). The AR also interacts with BRD4, and both full-length AR as well as constitutively-active splice variants can be inhibited via BETi (47, 48). Based on this work, multiple BETi are undergoing clinical trial evaluation, with evidence of clinical activity in hematologic malignancies (17). Furthermore, BETi are also in early-phase trials in mCRPC patients (NCT02705469, NCT02711956, and NCT04471974), where they have shown encouraging results when combined with AR antagonists (49). This is further supported by our data demonstrating that concurrent ADT enhances the anti-tumor efficacy of BETi and checkpoint blockade, suggesting that this approach may be useful in the neoadjuvant hormone-sensitive setting in patients with high-risk PC. Given the mixed results from single-agent trials utilizing BET inhibitors in most solid tumors, where pharmacodynamic inhibition does not consistently correlate with clinical responsiveness (50), the combination of ADT/BETi could enhance both tumor cell intrinsic cell death and extrinsic immune-based anti-cancer activity. Our data suggests that utilizing precision medicine for patient selection and immune-oncology combinations provide a means to amplify the tumor cell-extrinsic effects of BETi, thereby improving anti-tumor efficacy. Furthermore, the combination approach with ADT is aimed at pharmacological inhibition of BET proteins with biologically effective doses to amplify immunological activation and not the high doses needed to elicit tumor cell death, with the potential of avoiding toxicities associated with maximum tolerated doses in BETi clinical trials. In summary, the findings in this study provide a “proof-of-mechanism” for clinical translation of BET inhibitors with ICB to treat Rb-deficient cancers, including aggressive-variant Rb-deficient mCRPC.

## ACKNOWLEDGEMENTS

The results shown here are in whole or part based upon data generated by the TCGA Research Network (https://www.caner.gov/tcga).

## AUTHORS CONTRIBUTION

BMO designed and performed experiments, analyzed data, provided overall supervision for the project, and wrote the manuscript. KC designed experiments, performed experiments, analyzed data, and edited the manuscript. RB and SSS performed bioinformatic analysis and edited the manuscript. CJH, SR, RC, and AJT performed experiments. AP provided overall supervision for the project, designed experiments, analyzed data, and wrote the manuscript.

## SUPPLEMENTARY DATA

**Supplementary Table 1.** List of antibodies used in experiments.

| **Antibody** | **Catalog** | **Source** | **Application** | **Concentration** |
| --- | --- | --- | --- | --- |
| NKG2D | 563694 | BD Bioscience | FC | 1:100 |
| PD-L1 | 124314 | BioLegend | FC | 1:100 |
| CD45 | 103138 | BioLegend | FC | 1:100 |
| CD3 | 100306 | BioLegend | FC | 1:100 |
| mCD8 | 100750 | BioLegend | FC | 1:100 |
| hCD8 | 301010 | BioLegend | FC | 1:100 |
| CD4 | 100548 | BioLegend | FC | 1:100 |
| PD-1 | 135228 | BioLegend | FC | 1:100 |
| Tim3 | 119723 | BioLegend | FC | 1:100 |
| CD137 | 106106 | BioLegend | FC | 1:100 |
| Ki67 | 652419 | BioLegend | FC | 1:100 |
| CD11b | 101257 | BioLegend | FC | 1:100 |
| CD11c | 117334 | BioLegend | FC | 1:100 |
| Ly6C | 128044 | BioLegend | FC | 1:100 |
| I-A/I-E | 107632 | BioLegend | FC | 1:100 |
| CD103 | 121485 | BioLegend | FC | 1:100 |
| CD206 | 141723 | Biolegend | FC | 1:100 |
| CXCR5 | 145517 | BioLegend | FC | 1:100 |
| Arg1 | IC5868A | R&D Systems | FC | 1:100 |
| CD19 | 65-0193-U100 | Tonbo Biosciences | FC | 1:100 |
| Foxp3 | 20-0191-U100 | Tonbo Biosciences | FC | 1:100 |
| Ly6G | 80-5931-U100 | Tonbo Biosciences | FC | 1:100 |
| CD24 | 50-0242-U100 | Tonbo Biosciences | FC | 1:100 |
| F4/80 | 65-4801-U100 | Tonbo Biosciences | FC | 1:100 |
| Rb | 9313S | Cell Signaling Technology | WB | 1:500 |
| β-actin | 4967S | Cell Signaling Technology | WB | 1:5000 |
| HEXIM-1 | Ab25388 | Abcam | WB | 1:10,000 |
| STING | 13647S | Cell Signaling Technology | WB | 1:1000 |
| cGAS | 31659S | Cell Signaling Technology | WB | 1:1000 |
| Phospho-histone H2AX | 9718S | Cell Signaling Technology | WB | 1:1000 |
| Phospho-IRF3 (Ser396) | 29047 | Cell Signaling Technology | WB and ImageStream | 1:1000 (WB), 1:100 (ImageStream) |
| IRF3 | 4302S | Cell Signaling Technology | WB | 1:1000 |
| Phospho-TBK1 (Ser172) | 5483 | Cell Signaling Technology | WB | 1:1000 |
| TBK1 | 3504 | Cell Signaling Technology | WB | 1:1000 |
| Phospho-NF-κB p65<br>(Ser536) | 3033 | Cell Signaling Technology | WB | 1:1000 |
| Phospho-NF-κB p65<br>(Ser536) | 3036 | Cell Signaling Technology | ImageStream | 1:100 |
| NF-κB | 8242 | Cell Signaling Technology | WB | 1:1000 |
| Donkey anti-<br>rabbit IgG | 406414 | BioLegend | ImageStream | 1:200 |
| Goat anti-mouse | 405307 | BioLegend | ImageStream | 1:200 |
| Anti-PD-L1 | BE0101 | BioXCell | *in vivo* | 200µg/treatment |
| Isotype | BE0090 | BioXCell | *in vivo* | 200µg/treatment |
| Anti-CD4 | BE0003-1 | BioXCell | *in vivo* | 200µg/treatment |
| Anti-CD8 | BE0061 | BioXCell | *in vivo* | 200µg/treatment |

**Supplementary Table 2.**
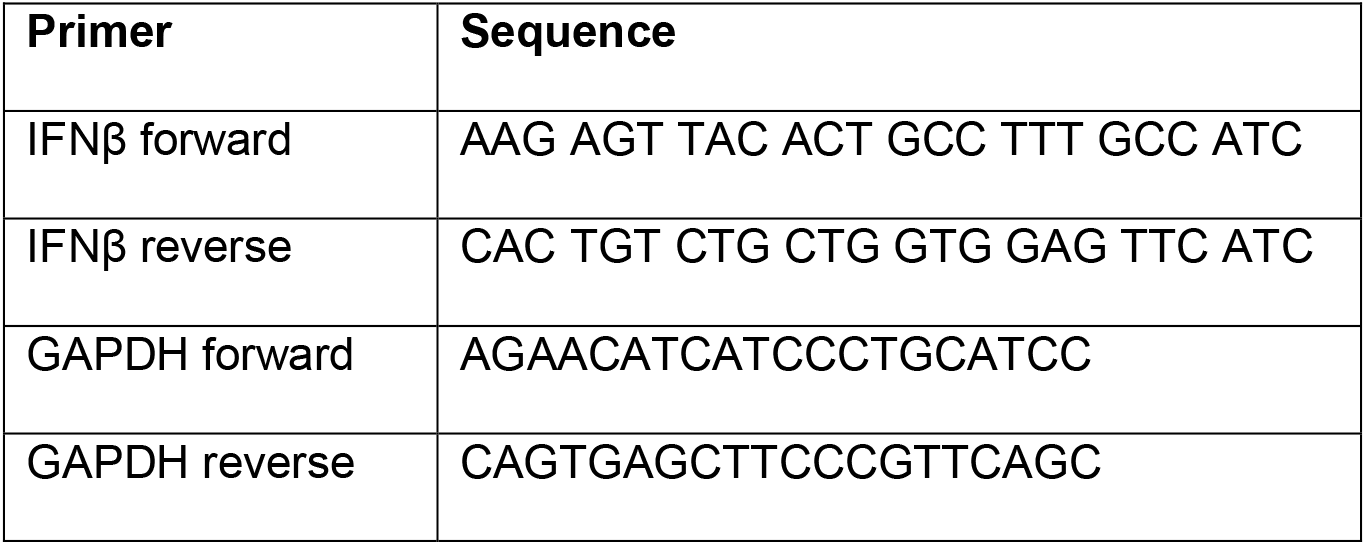
List of oligonucleotides.

**Supplementary Figure 1.**
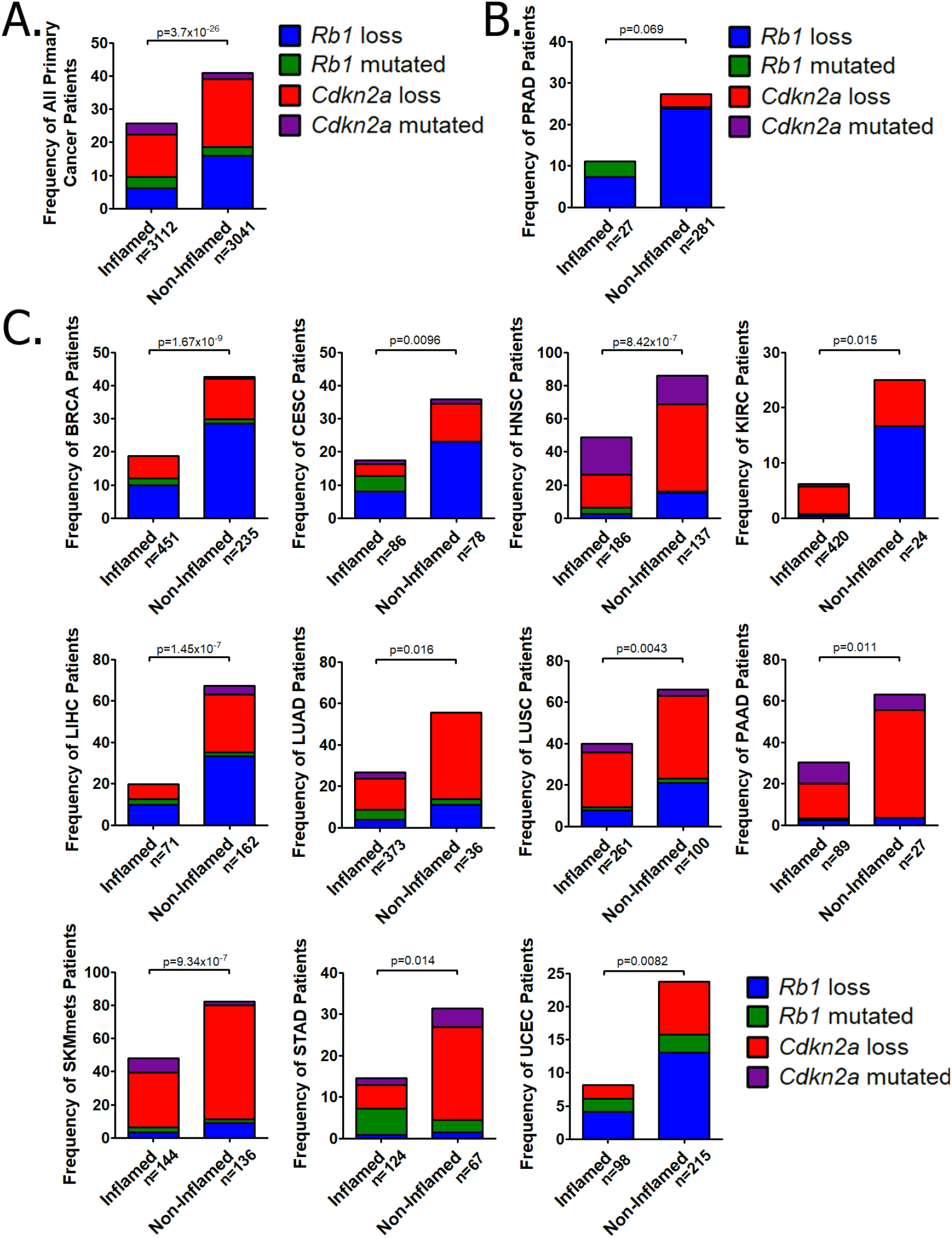
Enrichment of *Rb1* or *CDKN2A*loss in non-T cell-inflamed primary prostate cancer and across multiple primary tumor histologies. Primary tumors of all histologies (***panel A***), prostate cancer (***panel B***), or other histologies (***panel C***) on the TCGA database were grouped into T cell-inflamed or non-T cell-inflamed tumors, and exomic and RNAseq data was used to quantify the frequency of patients with loss of *Rb1*(*blue*), *Cdkn2a* (red), *Rb1* LOF mutations (*green*), or *Cdkn2a* LOF mutations (*purple*). The following abbreviations were used to indicate tumor types: BRCA: Breast invasive carcinoma; CESC: Cervical squamous cell carcinoma; HNSC: Head and Neck squamous cell carcinoma; KIRC: Kidney renal clear cell carcinoma; LIHC: Liver hepatocellular carcinoma; LUAD: Lung adenocarcinoma; LUSC: Lung squamous cell carcinoma; PAAD: Pancreatic adenocarcinoma; PRAD: Prostate adenocarcinoma; SKCM: Skin Cutaneous Melanoma; STAD: Stomach adenocarcinoma; UCEC: Uterine Corpus Endometrial Carcinoma.

**Supplementary Figure 2.**
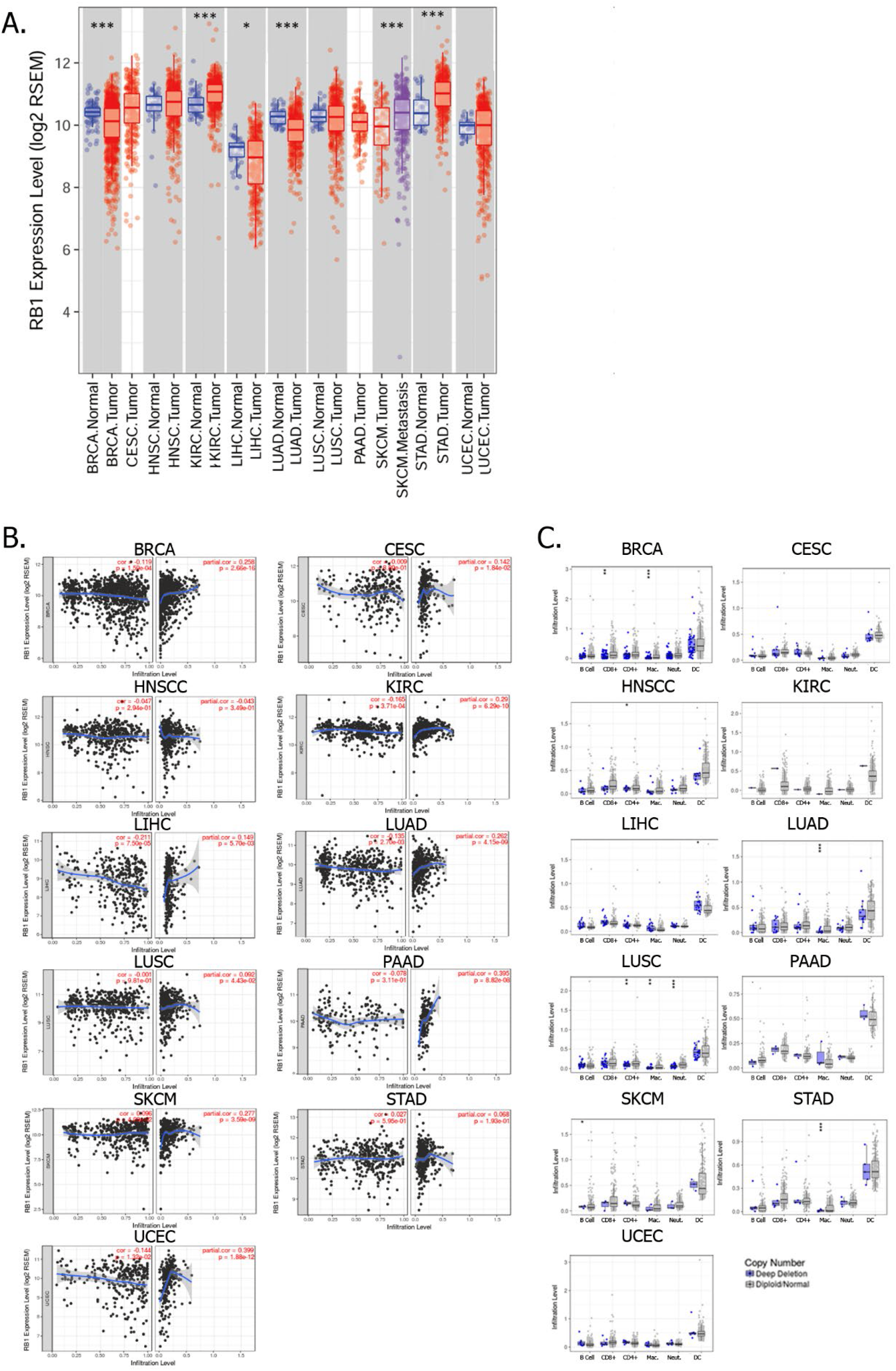
Decreased *Rb1* expression correlates with predicted decreased immune infiltration across multiple disease histologies. ***Panel A***, Gene expression data from primary tumors along with the adjacent normal tissue (when available) in the TCGA database was utilized for the estimation of expression profile of *Rb1* across cancer types. ***Panel B***, Gene expression level of *Rb1* was compared with estimated abundance tumor cells (to measure tumor purity; *left panels*) and CD8 T cells (*right panels*) using TIMER. ***Panel C***, Somatic Copy Number Alteration (SCNA) data from individual histologies in the TCGA database was utilized for the estimation of copy number status of *Rb1* (either deep deletion or normal diploid, as defined by GISTIC2.0(29)), and were analyzed for estimated B cell, CD4+ and CD8+ T cell, macrophage (Mac.), neutrophil (Neut.), and dendritic cell (DC) frequencies using TIMER. *: p < 0.05; **: p-value <0.01; ***: p<0.001, all by Wilcoxin signed-rank test.

**Supplementary Figure 3.**
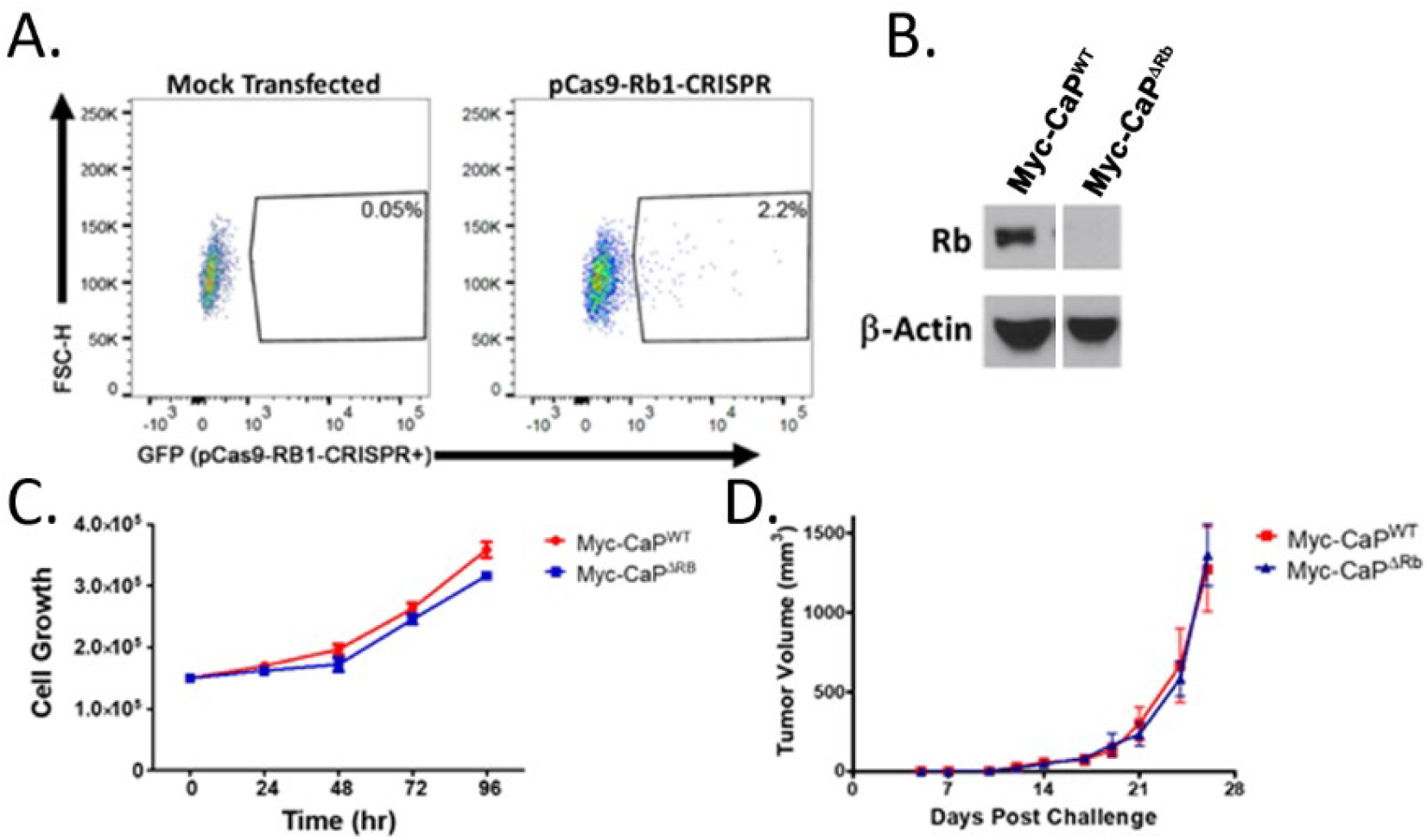
Generation and characterization of growth kinetics of Rb-deficient Myc-CaP cell line. ***Panel A***, Myc-CaP tumor cells were transfected with a pCas9-Rb1-CRISPR plasmid (or a mock control), and 24 hours later transfected cells were identified and single-cell sorted via FACS. ***Panel B***, Clonal cell lines were evaluated for Rb1 expression via Western blotting. ***Panel C,*** Myc-CaP^WT^ or Myc-CaP^ΔRb^ cells were characterized for growth kinetics *in vitro* using Trypan B exclusion. ***Panel D***, wild-type male FVB mice were challenged subcutaneously with Myc-CaP^WT^ or Myc-CaP^ΔRb^ tumor cells and followed for tumor growth.

**Supplementary Figure 4.**
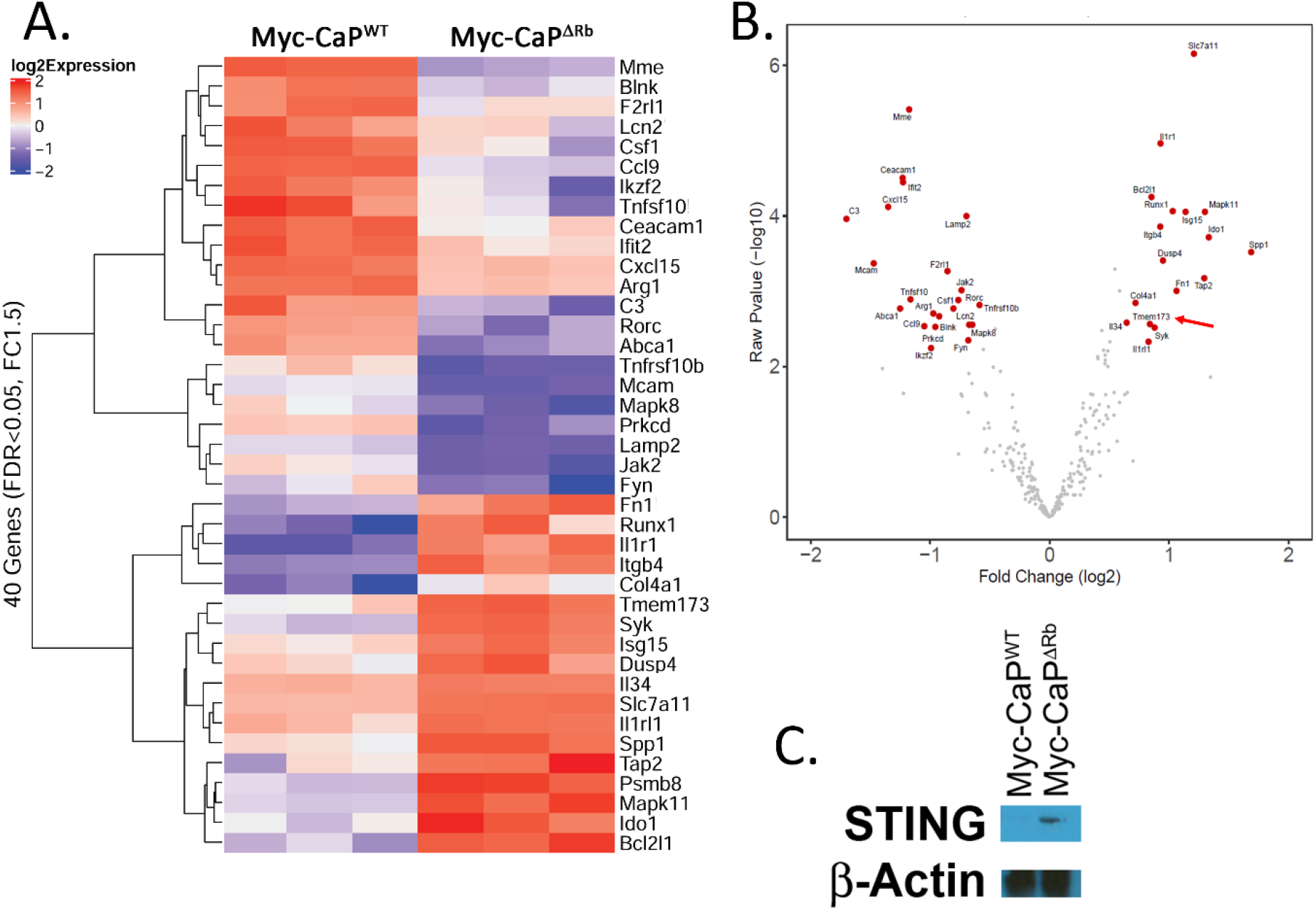
Loss of Rb results in altered gene expression that promotes immune suppression as well as increased *Tmem173/*STING expression. RNA from wild-type or Rb-deficient Myc-CaP cells were evaluated for the expression of various immune-related markers using the NanoString PanCancer Immune Profiling sequencing, with individual replicates shown on heat maps (***panel A***) and significant alterations identified in the Volcano plots (***panel B***), with *Tmem173* highlighted by an arrow in red. ***Panel C***, lysates from wild-type or Rb-deficient Myc-CaP cells were evaluated for STING expression by Western blot.

**Supplementary Figure 5.**
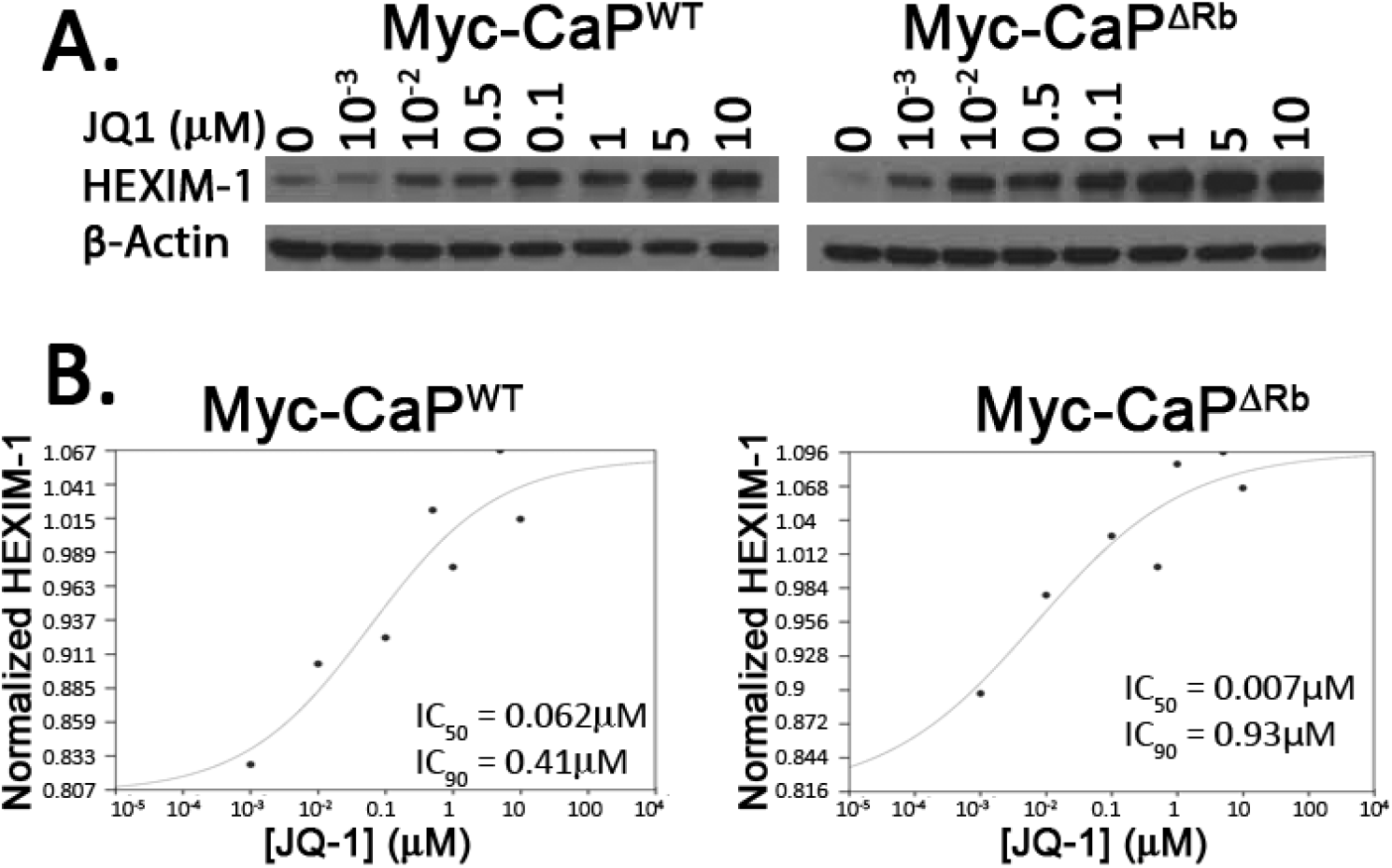
Pharmacological inhibition of BET family members by JQ-1. ***Panel A,*** Myc-CaP^WT^ or Myc-CaP^ΔRb^ cells were treated with increasing doses of JQ-1 for 24 hours and evaluated for pharmacodynamic inhibition of BRD family members via increased expression of HEXIM-1 by Western blot. This data was quantified via densitometry and used to calculate IC50 and IC90 using Quest Graph software (30) (***panel B***).

**Supplementary Figure 6.**
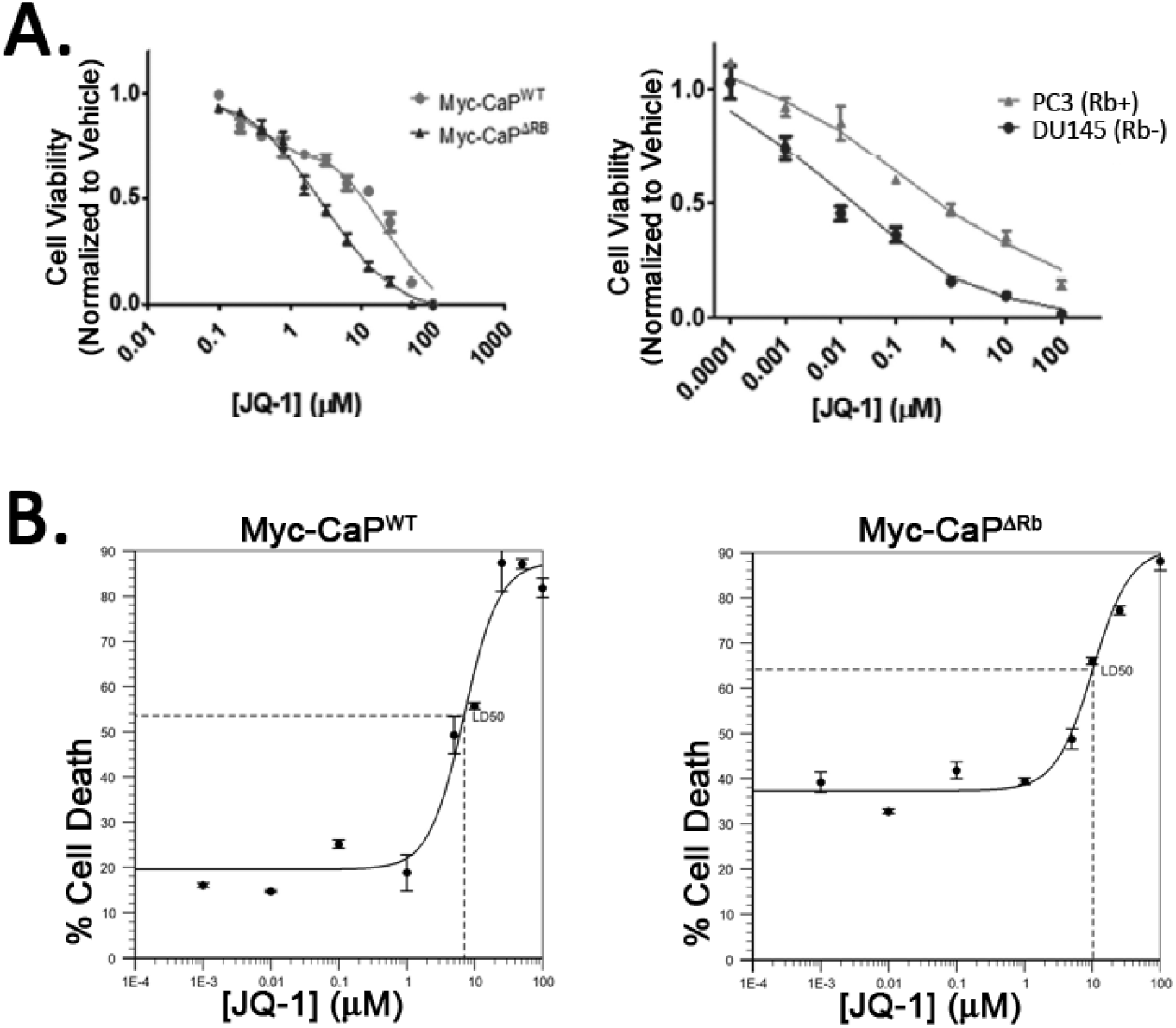
JQ-1 does not induce apoptosis at 1 µM concentration that activates non-canonical STING/NF-κB signaling. ***Panel A,*** Myc-CaP^WT^ or Myc-CaP^ΔRb^ (*left panels*) or human PC3 (Rb wild-type) or DU-145 (Rb-deficient) human prostate cancer cells (*right panels*) were treated with increasing doses of JQ-1 for 72 hours and evaluated for cell proliferation by Trypan B dye exclusion. ***Panel B***, to evaluate effects of BET inhibition on apoptosis, Myc-CaP^WT^ or Myc-CaP^ΔRb^ cells were treated with increasing doses of JQ-1 for 72 hours and evaluated for cell death via Annexin V-Propidium Iodide staining and flow cytometry. The percent dead/dying cells were calculated and used to calculate the LD50 using Quest Graph software (30).

**Supplementary Figure 7.**
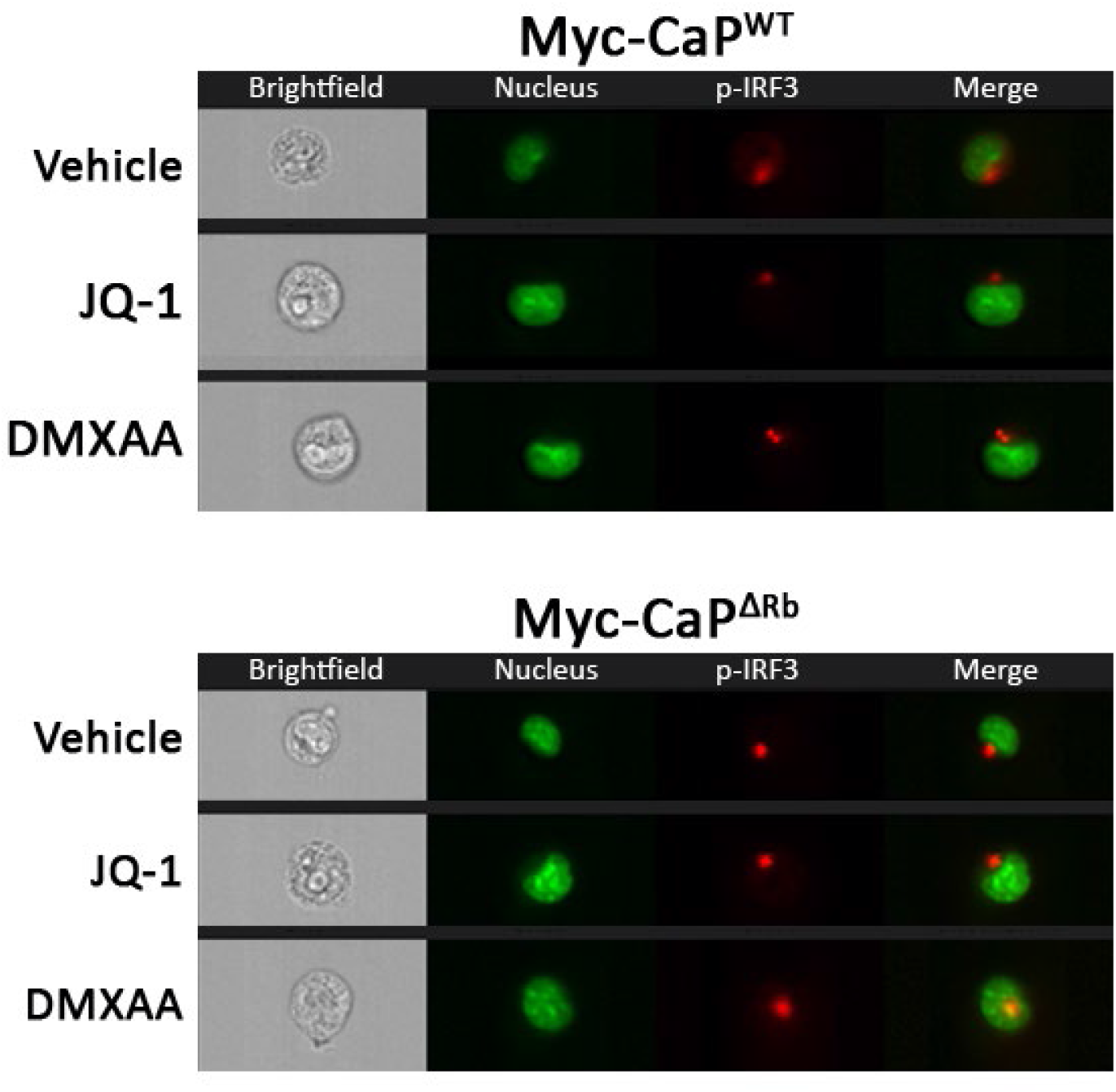
JQ-1 does not induce p-IRF3 translocation to the nucleus in wild-type or Rb-deficient Myc-CaP tumor cells. Myc-CaP^WT^ *(top panels)* or Myc-CaP^ΔRb^ (*bottom panels)* cells were treated with 1μM JQ-1 (or vehicle or DMXAA controls) for one hour, and nuclear expression of p-IRF3 was measured via ImageStream analysis. Panels demonstrate brightfield images, nuclear staining (green), p-IRF3 (red), or merged images.

**Supplementary Figure 8.**
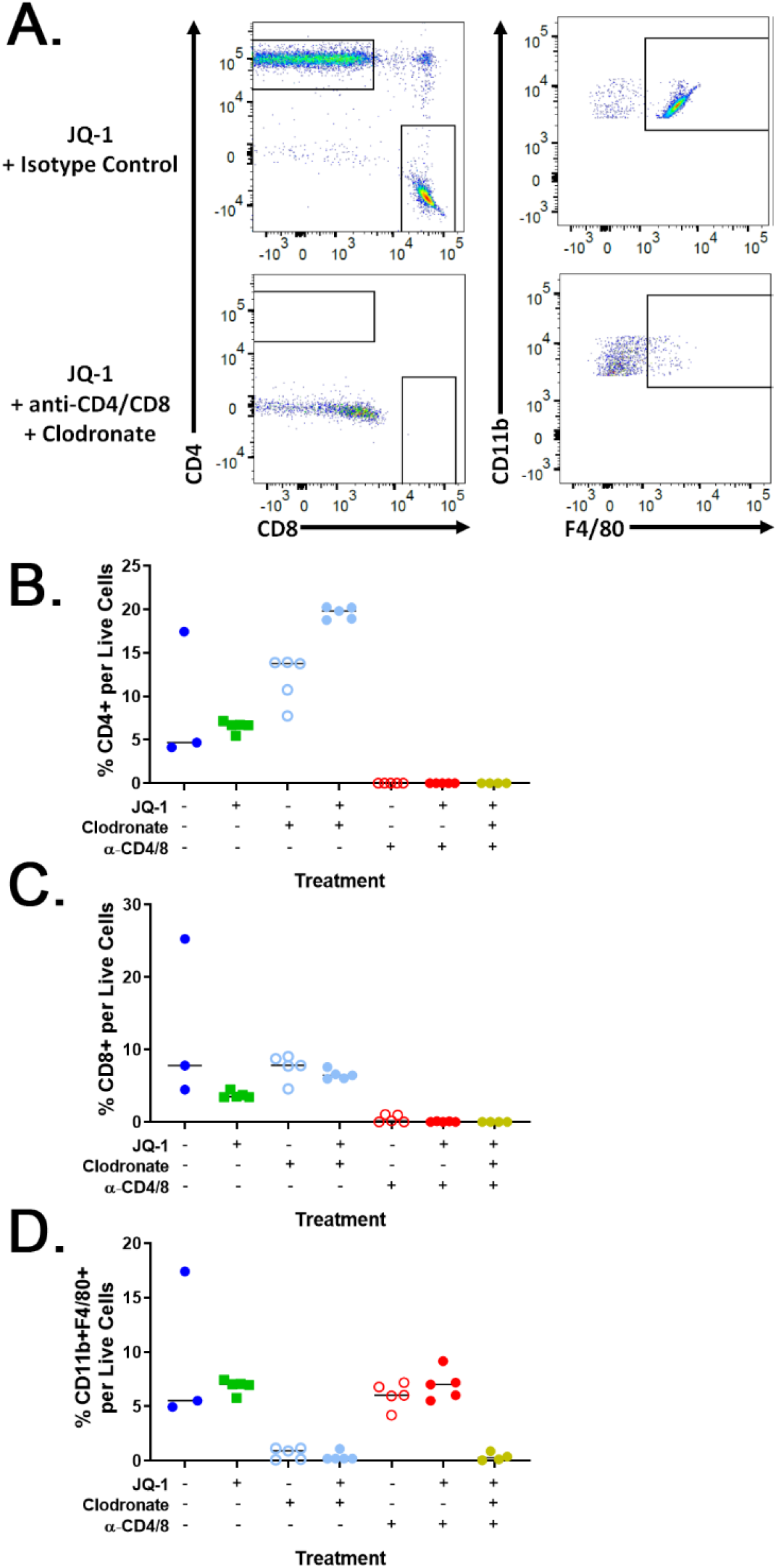
T cell and macrophage depletion studies in mice bearing Myc-CaP^WT^ or Myc-CaP^ΔRb^ tumors. Wild-type FVB mice bearing Myc-CaP^ΔRb^ tumors were treated with vehicle or JQ-1, either alone or combined with clodronate and/or anti-CD4/8 depletion antibodies. Spleens were collected and analyzed for efficacy of macrophage and T cell depletion by flow cytometry. ***Panel A***, Representative flow plot of splenocytes from a mouse treated with JQ-1 (*top panels*) or JQ-1+clodronate+anti-CD4/8 (*bottom panels*), stained for CD4 and CD8 T cells (*left panels*, gating on live/ dump-/CD45+ events) and macrophages (*right panels*, gating on live/dump-/CD45+/CD11b+ events). Frequency of cells were then quantified: CD4+ T cells (***panel B***), CD8+ T cells (***panel C***), or macrophages (***panel D***).

**Supplementary Figure 9.**
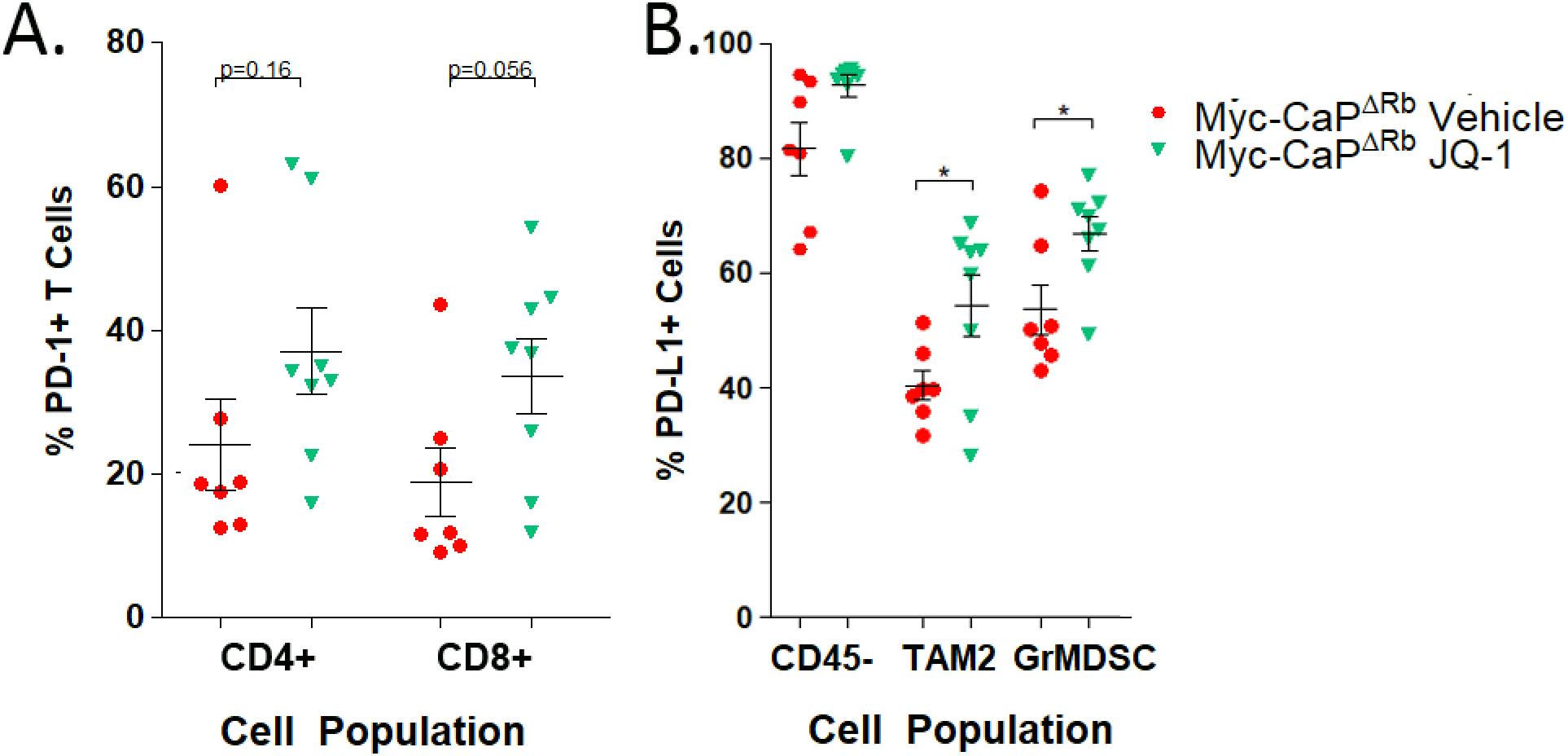
JQ-1 treatment *in vivo* increases PD-L1 expression on myeloid suppressive cells. Wild-type FVB mice were engrafted with Myc-CaP^ΔRb^ cells, and tumors were allowed to establish to 50mm^3^. Animals were treated for seven days with vehicle or JQ-1, and tumors were collected. Tumor-infiltrating CD4+ and CD8+ T cells (***panel A***), and CD45-cells and myeloid suppressive populations (***panel B***) were analyzed by flow cytometry for PD-1 expression and PD-L1 expression, respectively. In all panels, * indicates p<0.05 by student’s t test.

## REFERENCES

1. Pages F, Galon J, Dieu-Nosjean MC, Tartour E, Sautes-Fridman C, Fridman WH. Immune infiltration in human tumors: a prognostic factor that should not be ignored. Oncogene 2010;29(8):1093–102.

2. Spranger S, Bao R, Gajewski TF. Melanoma-intrinsic beta-catenin signalling prevents anti-tumour immunity. Nature 2015;523(7559):231–5 doi 10.1038/nature14404.

3. Ji RR, Chasalow SD, Wang L, Hamid O, Schmidt H, Cogswell J, et al. An immune-active tumor microenvironment favors clinical response to ipilimumab. Cancer Immunol Immunother 2011 doi 10.1007/s00262-011-1172-6.

4. Gajewski TF, Louahed J, Brichard VG. Gene signature in melanoma associated with clinical activity: a potential clue to unlock cancer immunotherapy. Cancer J 2010;16(4):399–403 doi 10.1097/PPO.0b013e3181eacbd8.

5. Robinson DR, Wu YM, Lonigro RJ, Vats P, Cobain E, Everett J, et al. Integrative clinical genomics of metastatic cancer. Nature 2017;548(7667):297–303 doi 10.1038/nature23306.

6. Burkhart DL, Sage J. Cellular mechanisms of tumour suppression by the retinoblastoma gene. Nat Rev Cancer 2008;8(9):671–82 doi 10.1038/nrc2399.

7. Taylor BS, Schultz N, Hieronymus H, Gopalan A, Xiao Y, Carver BS, et al. Integrative genomic profiling of human prostate cancer. Cancer cell 2010;18(1):11–22 doi 10.1016/j.ccr.2010.05.026.

8. Sharma A, Yeow WS, Ertel A, Coleman I, Clegg N, Thangavel C, et al. The retinoblastoma tumor suppressor controls androgen signaling and human prostate cancer progression. J Clin Invest 2010;120(12):4478–92 doi 10.1172/JCI44239.

9. Tan HL, Sood A, Rahimi HA, Wang W, Gupta N, Hicks J, et al. Rb loss is characteristic of prostatic small cell neuroendocrine carcinoma. Clin Cancer Res 2014;20(4):890–903 doi 10.1158/1078-0432.CCR-13-1982.

10. Ku SY, Rosario S, Wang Y, Mu P, Seshadri M, Goodrich ZW, et al. Rb1 and Trp53 cooperate to suppress prostate cancer lineage plasticity, metastasis, and antiandrogen resistance. Science 2017;355(6320):78–83 doi 10.1126/science.aah4199.

11. Markey MP, Bergseid J, Bosco EE, Stengel K, Xu H, Mayhew CN, et al. Loss of the retinoblastoma tumor suppressor: differential action on transcriptional programs related to cell cycle control and immune function. Oncogene 2007;26(43):6307–18 doi 10.1038/sj.onc.1210450.

12. Osborne AR, Zhang H, Blanck G. Oct-1 DNA binding activity unresponsive to retinoblastoma protein expression prevents MHC class II induction in a non-small cell lung carcinoma cell line. Mol Immunol 2006;43(6):710–6 doi 10.1016/j.molimm.2005.03.013.

13. Jin X, Ding D, Yan Y, Li H, Wang B, Ma L, et al. Phosphorylated RB Promotes Cancer Immunity by Inhibiting NF-kappaB Activation and PD-L1 Expression. Molecular cell 2019;73(1):22–35 e6 doi 10.1016/j.molcel.2018.10.034.

14. Hutcheson J, Bourgo RJ, Balaji U, Ertel A, Witkiewicz AK, Knudsen ES. Retinoblastoma protein potentiates the innate immune response in hepatocytes: significance for hepatocellular carcinoma. Hepatology 2014;60(4):1231–40 doi 10.1002/hep.27217.

15. Vidotto T, Nersesian S, Graham C, Siemens DR, Koti M. DNA damage repair gene mutations and their association with tumor immune regulatory gene expression in muscle invasive bladder cancer subtypes. J Immunother Cancer 2019;7(1):148 doi 10.1186/s40425-019-0619-8.

16. Filippakopoulos P, Knapp S. Targeting bromodomains: epigenetic readers of lysine acetylation. Nat Rev Drug Discov 2014;13(5):337–56 doi 10.1038/nrd4286.

17. Fu LL, Tian M, Li X, Li JJ, Huang J, Ouyang L, et al. Inhibition of BET bromodomains as a therapeutic strategy for cancer drug discovery. Oncotarget 2015;6(8):5501–16 doi 10.18632/oncotarget.3551.

18. Shorstova T, Foulkes WD, Witcher M. Achieving clinical success with BET inhibitors as anti-cancer agents. Br J Cancer 2021;124(9):1478–90 doi 10.1038/s41416-021-01321-0.

19. Bhadury J, Nilsson LM, Muralidharan SV, Green LC, Li Z, Gesner EM, et al. BET and HDAC inhibitors induce similar genes and biological effects and synergize to kill in Myc-induced murine lymphoma. Proceedings of the National Academy of Sciences 2014;111(26):E2721–E30 doi doi:10.1073/pnas.1406722111.

20. Manzotti G, Ciarrocchi A, Sancisi V. Inhibition of BET Proteins and Histone Deacetylase (HDACs): Crossing Roads in Cancer Therapy. Cancers (Basel) 2019;11(3):304 doi 10.3390/cancers11030304.

21. Dalgard CL, Van Quill KR, O’Brien JM. Evaluation of the in vitro and in vivo antitumor activity of histone deacetylase inhibitors for the therapy of retinoblastoma. Clin Cancer Res 2008;14(10):3113–23 doi 10.1158/1078-0432.CCR-07-4836.

22. Poulaki V, Mitsiades CS, Kotoula V, Negri J, McMullan C, Miller JW, et al. Molecular sequelae of histone deacetylase inhibition in human retinoblastoma cell lines: clinical implications. Invest Ophthalmol Vis Sci 2009;50(9):4072–9 doi 10.1167/iovs.09-3517.

23. Wang M, Zhao L, Tong D, Yang L, Zhu H, Li Q, et al. BET bromodomain inhibitor JQ1 promotes immunogenic cell death in tongue squamous cell carcinoma. Int Immunopharmacol 2019;76:105921 doi 10.1016/j.intimp.2019.105921.

24. Zhu H, Bengsch F, Svoronos N, Rutkowski MR, Bitler BG, Allegrezza MJ, et al. BET Bromodomain Inhibition Promotes Anti-tumor Immunity by Suppressing PD-L1 Expression. Cell Rep 2016;16(11):2829–37 doi 10.1016/j.celrep.2016.08.032.

25. Hogg SJ, Vervoort SJ, Deswal S, Ott CJ, Li J, Cluse LA, et al. BET-Bromodomain Inhibitors Engage the Host Immune System and Regulate Expression of the Immune Checkpoint Ligand PD-L1. Cell Rep 2017;18(9):2162–74 doi 10.1016/j.celrep.2017.02.011.

26. Mao W, Ghasemzadeh A, Freeman ZT, Obradovic A, Chaimowitz MG, Nirschl TR, et al. Immunogenicity of prostate cancer is augmented by BET bromodomain inhibition. J Immunother Cancer 2019;7(1):277 doi 10.1186/s40425-019-0758-y.

27. Kagoya Y, Nakatsugawa M, Yamashita Y, Ochi T, Guo T, Anczurowski M, et al. BET bromodomain inhibition enhances T cell persistence and function in adoptive immunotherapy models. J Clin Invest 2016;126(9):3479–94 doi 10.1172/JCI86437.

28. Luke JJ, Bao R, Sweis RF, Spranger S, Gajewski TF. WNT/beta-catenin Pathway Activation Correlates with Immune Exclusion across Human Cancers. Clin Cancer Res 2019;25(10):3074–83 doi 10.1158/1078-0432.CCR-18-1942.

29. Mermel CH, Schumacher SE, Hill B, Meyerson ML, Beroukhim R, Getz G. GISTIC2.0 facilitates sensitive and confident localization of the targets of focal somatic copy-number alteration in human cancers. Genome Biol 2011;12(4):R41 doi 10.1186/gb-2011-12-4-r41.

30. AAT Bioquest I. 2021 May 30, 2021. Quest Graph™ LD50 Calculator (v.1). <https://www.aatbio.com/tools/ld50-calculator-v1 > May 30, 2021.

31. Robinson D, Van Allen EM, Wu YM, Schultz N, Lonigro RJ, Mosquera JM, et al. Integrative clinical genomics of advanced prostate cancer. Cell 2015;161(5):1215–28 doi 10.1016/j.cell.2015.05.001.

32. Watson PA, Ellwood-Yen K, King JC, Wongvipat J, Lebeau MM, Sawyers CL. Context-dependent hormone-refractory progression revealed through characterization of a novel murine prostate cancer cell line. Cancer Res 2005;65(24):11565–71 doi 10.1158/0008-5472.CAN-05-3441.

33. Muralidharan SV, Bhadury J, Nilsson LM, Green LC, McLure KG, Nilsson JA. BET bromodomain inhibitors synergize with ATR inhibitors to induce DNA damage, apoptosis, senescence-associated secretory pathway and ER stress in Myc-induced lymphoma cells. Oncogene 2016;35(36):4689–97 doi 10.1038/onc.2015.521.

34. Filippakopoulos P, Qi J, Picaud S, Shen Y, Smith WB, Fedorov O, et al. Selective inhibition of BET bromodomains. Nature 2010;468(7327):1067–73 doi 10.1038/nature09504.

35. Olson BM, Gamat M, Seliski J, Sawicki T, Jeffery J, Ellis L, et al. Prostate Cancer Cells Express More Androgen Receptor (AR) Following Androgen Deprivation, Improving Recognition by AR-Specific T Cells. Cancer Immunol Res 2017;5(12):1074–85 doi 10.1158/2326-6066.CIR-16-0390.

36. Berchuck JE, Viscuse PV, Beltran H, Aparicio A. Clinical considerations for the management of androgen indifferent prostate cancer. Prostate Cancer Prostatic Dis 2021;24(3):623–37 doi 10.1038/s41391-021-00332-5.

37. Witkiewicz AK, Chung S, Brough R, Vail P, Franco J, Lord CJ, et al. Targeting the Vulnerability of RB Tumor Suppressor Loss in Triple-Negative Breast Cancer. Cell Rep 2018;22(5):1185–99 doi 10.1016/j.celrep.2018.01.022.

38. Lee C, Kim JK. Chromatin regulators in retinoblastoma: Biological roles and therapeutic applications. J Cell Physiol 2021;236(4):2318–32 doi 10.1002/jcp.30022.

39. Peng W, Chen JQ, Liu C, Malu S, Creasy C, Tetzlaff MT, et al. Loss of PTEN Promotes Resistance to T Cell-Mediated Immunotherapy. Cancer Discov 2016;6(2):202–16 doi 10.1158/2159-8290.cd-15-0283.

40. Bosco EE, Mayhew CN, Hennigan RF, Sage J, Jacks T, Knudsen ES. RB signaling prevents replication-dependent DNA double-strand breaks following genotoxic insult. Nucleic Acids Res 2004;32(1):25–34 doi 10.1093/nar/gkg919.

41. Lam FC, Kong YW, Huang Q, Vu Han TL, Maffa AD, Kasper EM, et al. BRD4 prevents the accumulation of R-loops and protects against transcription-replication collision events and DNA damage. Nat Commun 2020;11(1):4083 doi 10.1038/s41467-020-17503-y.

42. Zhang B, Lyu J, Liu Y, Wu C, Yang EJ, Pardeshi L, et al. BRCA1 deficiency sensitizes breast cancer cells to bromodomain and extra-terminal domain (BET) inhibition. Oncogene 2018;37(49):6341–56 doi 10.1038/s41388-018-0408-8.

43. Chakraborty G, Armenia J, Mazzu YZ, Nandakumar S, Stopsack KH, Atiq MO, et al. Significance of BRCA2 and RB1 Co-loss in Aggressive Prostate Cancer Progression. Clin Cancer Res 2019 doi 10.1158/1078-0432.CCR-19-1570.

44. Dunphy G, Flannery SM, Almine JF, Connolly DJ, Paulus C, Jonsson KL, et al. Non-canonical Activation of the DNA Sensing Adaptor STING by ATM and IFI16 Mediates NF-kappaB Signaling after Nuclear DNA Damage. Mol Cell 2018;71(5):745–60 e5 doi 10.1016/j.molcel.2018.07.034.

45. Deng G, Zeng F, Su J, Zhao S, Hu R, Zhu W, et al. BET inhibitor suppresses melanoma progression via the noncanonical NF-κB/SPP1 pathway. Theranostics 2020;10(25):11428–43 doi 10.7150/thno.47432.

46. Qiao J, Chen Y, Mi Y, Jin H, Wang L, Huang T, et al. Macrophages confer resistance to BET inhibition in triple-negative breast cancer by upregulating IKBKE. Biochem Pharmacol 2020;180:114126 doi 10.1016/j.bcp.2020.114126.

47. Asangani IA, Dommeti VL, Wang X, Malik R, Cieslik M, Yang R, et al. Therapeutic targeting of BET bromodomain proteins in castration-resistant prostate cancer. Nature 2014;510(7504):278–82 doi 10.1038/nature13229.

48. Welti J, Sharp A, Yuan W, Dolling D, Nava Rodrigues D, Figueiredo I, et al. Targeting Bromodomain and Extra-Terminal (BET) Family Proteins in Castration-Resistant Prostate Cancer (CRPC). Clin Cancer Res 2018;24(13):3149–62 doi 10.1158/1078-0432.CCR-17-3571.

49. Aggarwal R, Abida W, Schweizer M, Pantuck A, Nanus D, Heath E, et al. Abstract CT095: A Phase Ib/IIa study of the BET bromodomain inhibitor ZEN-3694 in combination with enzalutamide in patients with metastatic castration-resistant prostate cancer (mCRPC). Cancer Research 2019;79(13 Supplement):CT095-CT doi 10.1158/1538-7445.am2019-ct095.

50. Sun YL, Han J, Wang ZZ, Li XN, Sun YH, Hu ZB. Safety and Efficacy of Bromodomain and Extra-Terminal Inhibitors for the Treatment of Hematological Malignancies and Solid Tumors: A Systematic Study of Clinical Trials. Front Pharmacol 2021;11 doi ARTN62109310.3389/fphar.2020.621093.

